# Prior to Undergoing Epithelial to Mesenchymal Transition, Premalignant Cells Transiently Increase Tensile Forces at Cell-Cell and Cell-Matrix Adhesions

**DOI:** 10.1101/2025.07.05.663275

**Authors:** Lídia Faria, Carine M Gonçalves, Inês Fidalgo, Carla S. Lopes, Vanessa Rodrigues, Maria M. Azevedo, Paula Sampaio, Sandra Tavares, Vaibhav Mahajan, Anna Taubenberger, Florence Janody

## Abstract

The weakening of Adherens Junctions (AJs) and Focal Adhesions (FAs) adhesiveness has been proposed to enable the initiation and progression of several types of cancer. Here we report that, prior to disassembling AJs and acquiring malignant traits, premalignant mammary epithelial cells overactivating the Src proto-oncogene transiently enhance tensile forces at both AJs and FAs, thereby gaining a proliferative advantage. We show that AJs and FAs are transiently under higher tensile forces in premalignant Src-activated cells. Cells unable to increase tensile forces at FAs by knocking down PXN or inhibiting FAK activity fail to transiently build up tensile force at AJs and to grow. Conversely, preventing AJ strengthening using EGTA or small interference RNA against P-cadherin suppresses the transient increase in tensile forces at FAs, EGFR-ERK and MRTF-A-SRF activation and cell proliferation. Moreover, knocking down E-cadherin, the sole classical cadherin in *Drosophila*, inhibits Src-induce tissue overgrowth *in vivo*. Thus, prior to the loss of cell-cell and cell-matrix adhesiveness, strengthening of AJs and FAs may be an essential early step for mammary cells to gain a proliferative advantage and establish the mechanical and signaling conditions necessary for subsequent malignant transformation.

**Statement of Significance:** Although the weakening of cell-cell and cell-matrix adhesiveness is commonly associated with cancer initiation and progression, our findings reveal that an initial increase in adhesiveness enables proliferation and malignant progression.

## Introduction

Most breast cancers arise in the epithelial ducts and are referred to as ductal carcinoma. The most acknowledged model of ductal breast cancer progression states that every invasive carcinoma originates from a benign hyperplasia. These lesions are caused by excessive proliferation of the terminal duct lobular units, leading to dilated ducts, lined with one or more layers of epithelial cells (1). Progression of these proliferative breast lesions to invasive carcinoma has been proposed to involve a shift from an epithelial to a mesenchymal phenotype, in a process referred to as Epithelial to Mesenchymal Transition (EMT), characterized by the breakdown of Adherens Junctions (AJs) and dynamic remodeling of Focal Adhesions (FAs) (2).

AJs are the major cell-cell adhesion structure that senses and responds to tensile forces at the intercellular contact interface and hold epithelial cells together (3). AJs assemble through homotypic interactions between the extracellular domains of classical cadherins from adjacent cells in a Ca^2+^-dependent manner. E-cadherin (Ecad) is one of the main classical cadherins, highly conserved from bilaterian metazoans to mammals and expressed in most epithelial tissues (4). Intracellularly, the Ecad cytoplasmic tail interacts with p120-catenin (p120ctn) and β-catenin (βcat). While p120ctn promotes the assembly of Ecad dimers, βcat interacts with the actin-binding protein α-catenin (αcat) and physically links AJ complexes to the actomyosin cytoskeleton in a force-dependent manner (5). P-cadherin (Pcad), another classical cadherin, is also expressed in several epithelia, where it can overlap with Ecad. Alike Ecad, Pcad promotes cell-cell adhesion and Ca^2+^-dependent cell-cell aggregation (6).

At the cell-extracellular matrix (ECM) interface, mechanosensitive FAs assemble from nascent adhesions, which form upon the binding of a few integrin dimers to their ECM ligands and to Talin intracellularly (7). The coupling of integrins to the actomyosin cytoskeleton through Talin and Paxillin recruits Vinculin, leading to the maturation of nascent adhesions into larger and more stable FAs. The mechanisms in play involve the stretching of Talin and Vinculin by tensile forces, thereby re-enforcing the connection between integrins and the actomyosin cytoskeleton and the recruitment of additional proteins, including Focal Adhesion Kinase (FAK) and Zyxin to stress fiber-bound FAs (8,9). Thus, Ecad and Vinculin are *bona fide* mechanosensors (9,10).

In addition to maintaining cell–cell contact and attachment to the ECM, piconewton mechanical forces transmitted at AJs and FAs expose cryptic sites within mechanosensitive proteins and modulate the binding kinetics of receptor-ligand complexes to regulate the activity of diverse mechanotransduction signaling pathways and therefore cell behavior (11). Mechanical forces associated with FAs induce the nuclear translocation of the myocardin-related transcription factor (MRTF-A), a co-factor of serum response factor (SRF), which triggers invasion in mammary epithelial cells (12). Likewise, ECM-dependent FA strengthening promotes mammary epithelial cell growth through activation of extracellular signal-regulated kinase (ERK) (13). Increasing tension on homophilic Ecad bonds has also been shown to stimulate ERK-dependent proliferation by enabling the homodimerization of epidermal growth factor receptor (EGFR) in A-431D epithelial cells (14). Forces on AJs can also be transferred to FAs and *vice versa*. Integrin engagement or higher traction force from the ECM can increase tension at cell-cell contacts in multiple cell types (15,16). Conversely, Ecad force transmission increases the average FA area and number and Myosin-dependent cell contractility in MCF7 cells (17). In turn, integrins strengthen tensile forces at AJs, thus, establishing a positive mechanotransduction feedback loop between force-activated Ecad and integrins (18). However, strengthening of one adhesion can also antagonize the assembly of the other (11). How the balance of forces between adhesion sites controls distinct outcomes during cancer progression is not fully understood.

Using the mammary epithelial cells MCF10A-ER-Src (ER-Src), which contains a fusion between the non-receptor tyrosine kinase v-Src and the ligand-binding domain of the Estrogen Receptor (ER-Src), inducible with tamoxifen (TAM) treatment, we have reported that malignant progression involves two main phases. During the first 12 hours of Src activation, cells acquire a phenotypic state akin to premalignant cells, characterized by the gain of proliferative abilities, which depends on a transient increase in cell stiffening, and activation of ERK and MRTF-A/SRF signaling in cells that maintain an epithelial-like organization. 24 hours after Src activation, cells progress into a malignant phase, defined by the loss of epithelial-like organization associated with the downregulation of Ecad and Pcad and the upregulation of the EMT transcription factor SNAIL1 (19-21). We now provide evidence that the prior disassembling their AJs, premalignant ER-Src cells transiently increase tensile forces at AJs and FAs to acquire a proliferative advantage. Thus, tensile forces at AJs and FAs could have paradoxical roles during tumor progression. Strengthening of AJs and FAs may be an essential early step for premalignant breast cells to gain a proliferative advantage and establish the mechanical and signaling conditions necessary for subsequent malignant transformation. In contrast, the later weakening of forces at AJs and FAs could facilitate the emergence of malignant behaviors such as loss of adhesion, increased motility, and invasiveness.

## Results

### Src strengthens tensile forces at AJs prior to inducing EMT

Prior to acquiring malignant features, premalignant ER-Src cells turn transiently stiffer (19). We therefore asked if AJs were under higher tension in TAM-treated ER-Src cells. To this end, we generated ER-Src cells stably expressing an Ecad fluorescence resonance energy transfer (FRET)-based tension sensor (Ecad-TS), known to respond to changes in tensile forces at cell-cell contacts (10). The Ecad-TS was built by inserting a tension module (TS), containing a flagelliform elastic linker flanked by the mTFP1 donor and Venus acceptor fluorescent proteins in the Ecad coding sequence, between the transmembrane domain and the catenin-binding domain (Fig. 1A). As expected, the Ecad-TS was recruited at cell-cell contacts where it co-localized with p120ctn (Fig. 1B). To ensure that the Ecad-TS was functional, we tested if expressing the Ecad-TS in MDA-MB-231 cells that lack Ecad, restored cell-cell contacts. While p120ctn was barely detectable in control MDA-MB-231 cells, those expressing the Ecad-TS recruited p120ctn at the membrane between adjacent Ecad-TS expressing MDA-MB-231 cells and acquired a cobblestone-like shape (Fig. 1C). We next assessed if tensile forces applied to the Ecad-TS were altered during transformation of ER-Src cells by performing fluorescence lifetime measurement (FLIM) of the Ecad-TS donor, a quantitative imaging technique that is less sensitive to imaging or processing artifacts than the FRET ratio-metric approach (22). For FLIM analysis we focused on donor’s lifetimes, which increases as the distance with the acceptor increases. This approach revealed that the donor lifetime of the Ecad-TS was significantly extended in ER-Src cells treated with TAM for 6 or 12 hours, compared to those treated with the vehicle EtOH for the same time (Fig. 1D), suggesting that AJs are under higher tension in premalignant ER-Src cells. While cells undergo EMT 24 hours after TAM treatment (21), those that still maintained cell-cell contacts retained an extended donor lifetime of the Ecad-TS (Fig. 1D). To ensure that alterations of the Ecad-TS donor lifetime in TAM-treated cells resulted from higher tensile forces on Ecad, we also generated ER-Src cells expressing a control Ecad-TS construct lacking the βcat–binding domain (Ecad-TSΔcyto), which therefore cannot associate with F-actin (Fig. 1A). Ecad-TSΔcyto localized with p120ctn at cell-cell contacts in ER-Src cells (Fig. 1B), but did not recruit p120ctn at the cell membrane in MDA-MB-231 cells (Fig. 1C). The donor lifetime of the Ecad-TSΔcyto was not affected between EtOH and TAM-treated cells at any time point (Fig. 1E), indicating that the extended donor lifetime of the Ecad-TS in TAM-treated ER-Src cells depends on its ability to interact with the actin cytoskeleton. The Ecad-TS donor lifetime also depends on Myosin II activity, as Blebbistatin significantly lowered the Ecad-TS donor lifetime in cells treated with EtOH or TAM for 6 hours, compared to those grown in the absence of Blebbistatin (Supp. Fig. 1A).

**Figure 1:**
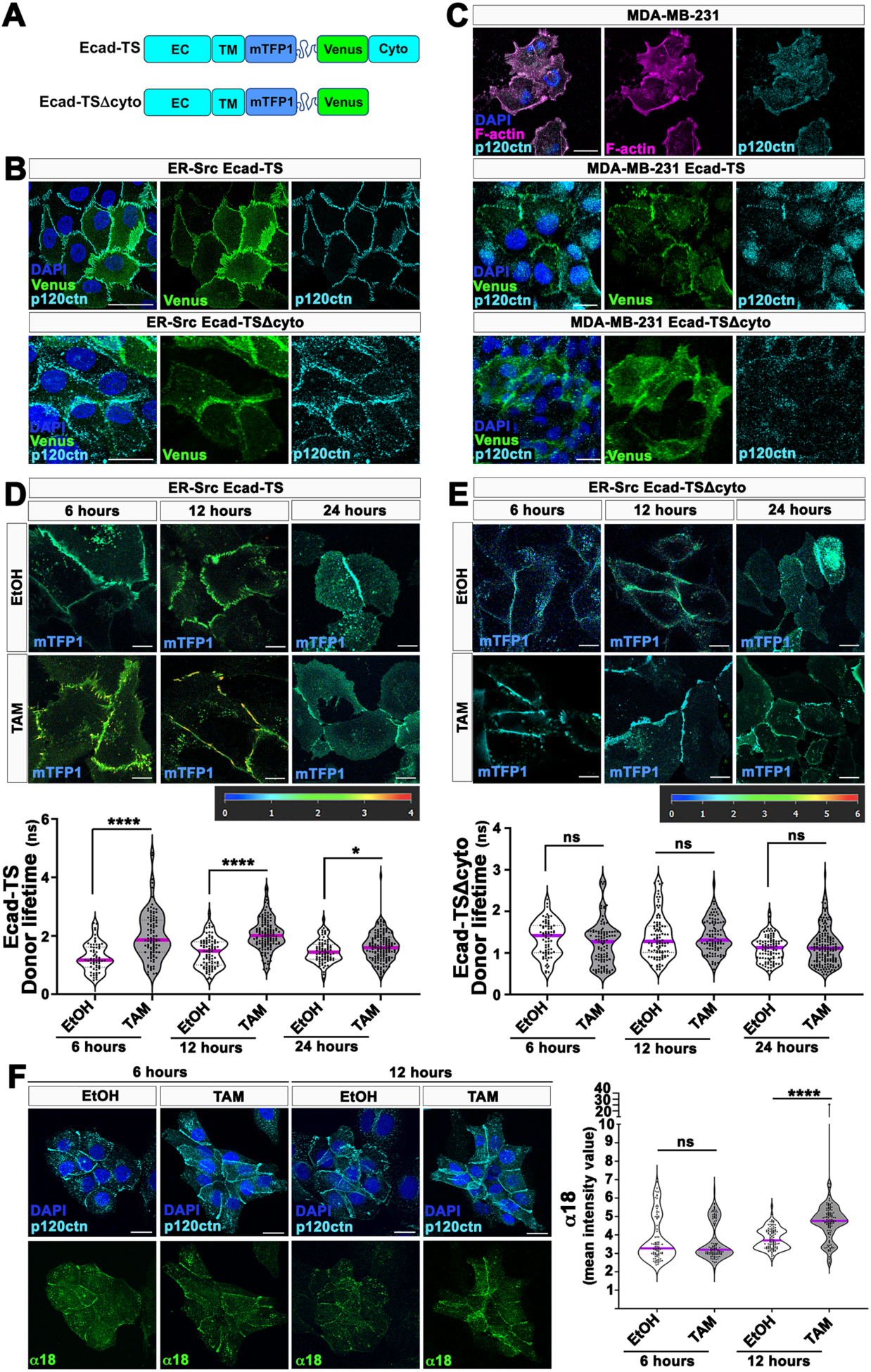
Prior to undergoing EMT, TAM-treated ER-Src cells strengthen tensile forces at AJs. **(A)** Schematics of the Ecad-TS and Ecad-TSΔcyto. EC: extracellular domain; TM transmembrane domain; cyto: cytoplasmic domain. **(B)** Confocal images of ER-Src cells expressing the Ecad-TS (Venus in green) or Ecad-TSΔcyto (Venus in green) and stained with anti-p120ctn (cyan) and DAPI (blue). **(C)** Confocal images of MDA-MB-231 cells, stained with anti-p120ctn (cyan), Phalloidin (magenta) and DAPI (blue) or expressing Ecad-TS (Venus in green) or Ecad-TSΔcyto (Venus in green), and stained with anti-p120ctn (cyan) and DAPI (blue). **(D)** (Top) ER-Src cells expressing Ecad-TS and treated with EtOH or TAM for 6, 12 or 24 hours. Color bar indicates 0–4 ns. (Bottom) Quantification of the mTFP1 lifetime at cell-cell contacts in ER-Src cells expressing the Ecad-TS and treated with EtOH or TAM for 6, 12 or 24 hours. **(E)** (Top) ER-Src cells expressing the Ecad-TSΔcyto and treated with EtOH or TAM for 6, 12 or 24 hours. Color bar indicates 0–6 ns. (Bottom) Quantification of the mTFP1 lifetime at cell-cell contacts in ER-Src cells expressing the Ecad-TSΔcyto and treated with EtOH or TAM for 6, 12 or 24 hours. Higher and lower donor lifetime correspond to higher and lower tension, respectively. **(F)** (Left) Confocal images of ER-Src cells treated with EtOH or TAM for 6 or 12 hours and stained with the α18 antibody (green), anti-p120ctn (cyan) and DAPI (blue). (Right) Quantification of the α18 mean intensity levels at cell-cell contacts in ER-Src cells treated with EtOH or TAM for 6 or 12 hours. Quantifications are from three biological replicates and are presented as violin plots with the magenta lines indicating median values. Statistical significance was calculated using Mann – Whitney t-tests. ns indicates non-significant. *P<0.05; ****P<0.0001. Scale bars represent 20 μm.

To confirm that prior to disassembling AJs 24 hours after TAM treatment, AJs are under higher tension, we stained cells with an antibody against the α18 epitope of αcat, a cryptic site revealed when tension is applied at AJs (23). While the intensity of the α18 antibody at cell-cell contacts was not significantly different between ER-Src cells treated with EtOH or TAM for 6 hours, it was significantly higher in cells treated with TAM for 12 hours, compared to those treated with EtOH for the same time (Fig. 1F). Because total αcat levels were not affected between EtOH and TAM-treated cells (Supp. Fig. 1B and C), we conclude that AJs are under higher tension in premalignant ER-Src cells, prior cells disassembling AJs.

### FAs are transiently under higher tension in premalignant ER-Src cells

Premalignant ER-Src cells accumulate polarized FA-associated stress fibers (19). We therefore asked if tension at FAs was also affected in premalignant ER-Src cells. We generated ER-Src cells expressing a Vinculin tension sensor (Vin-TS), which contains the tension module inserted between the head domain that recruits Vinculin to FAs and the tail domain, which binds to the mechanotransduction force machinery and responds to change in tension at FAs (9). To abrogate the ability of the Vin-TS to respond to tension, we also generated stable ER-Src cells expressing a Vin-TS lacking the tail domain (Vin-TSΔtail) (Fig. 2A). Vin-TS and Vin-TSΔtail mainly localized with Zyxin at FAs (Fig. 2B). FRET-FLIM analysis indicated that the donor lifetime of the Vin-TS at FAs was slightly extended 6 hours after TAM treatment, compared to those treated with EtOH for the same time. This effect was strongly enhanced 12 hours after TAM-treatment. However, 24 hours after treatment, this differential behavior was lost between EtOH- and TAM-treated cells (Fig. 2C). In contrast, the donor lifetime inVin-TSΔtail was not altered between EtOH- and TAM-treated cells at any time point (Fig. 2D). Moreover, Blebbistatin reduced the donor lifetime of the Vin-TS 12 hours after EtOH or TAM treatment (Supp. Fig. 2), indicating that the extended donor lifetime of the Vin-TS in TAM-treated ER-Src cells depends on Myosin II activity.

**Figure 2:**
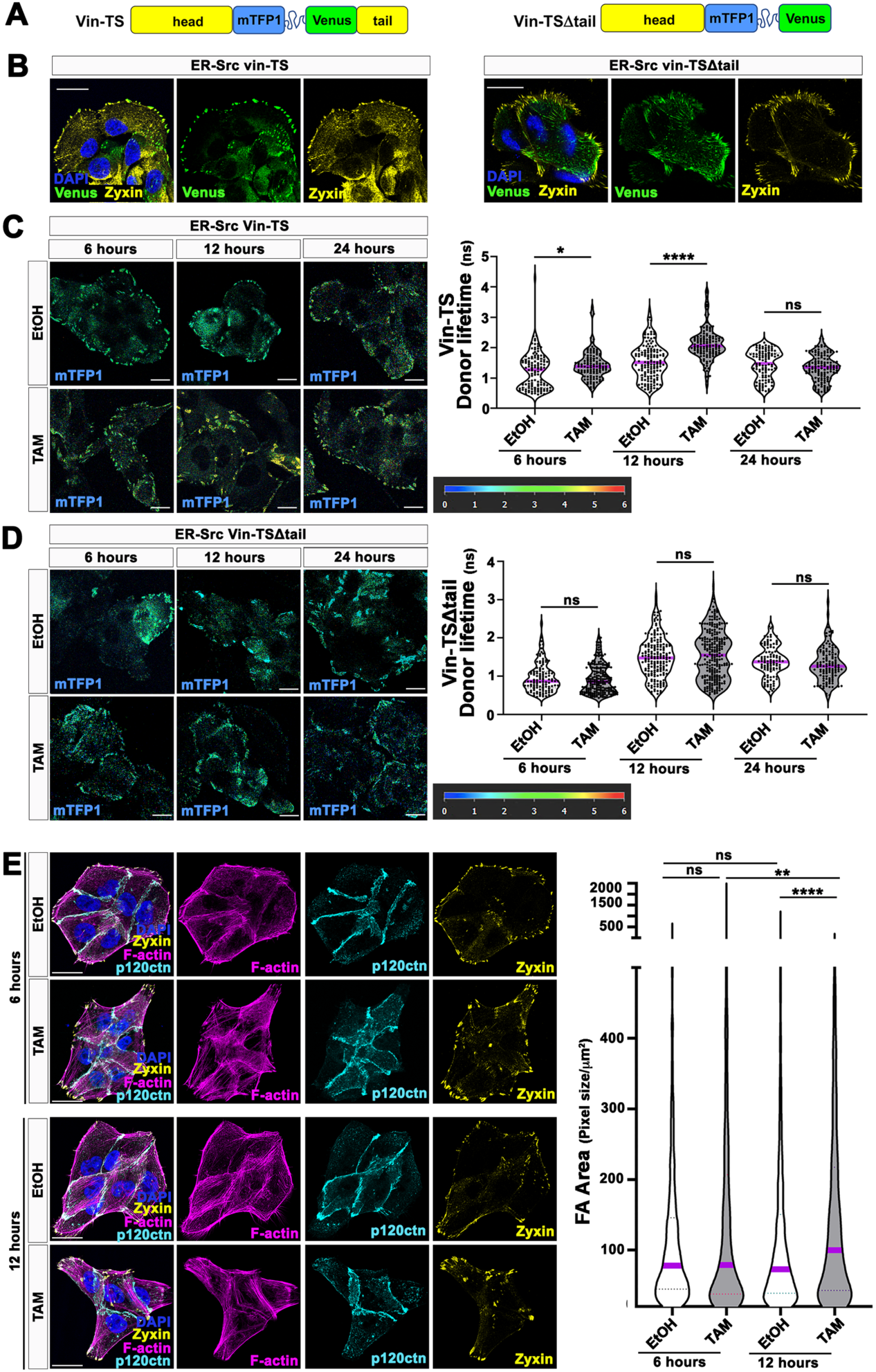
FAs are larger and transiently under higher tension in ER-Src cells. **(A)** Schematics of the Vin-TS and Vin-TSΔtail. **(B)** Confocal images of ER-Src cells expressing the Vin-TS (Venus in green) or Vin-TSΔtail (Venus in green) and stained with anti-Zyxin (yellow) and DAPI (blue). **(C)** (Left) ER-Src cells expressing the Vin-TS and treated with EtOH or TAM for 6, 12 or 24 hours. Color bar indicates 0–6 ns. (Right) Quantification of the mTFP1 lifetime at FAs in ER-Src cells expressing the Vin-TS and treated with EtOH or TAM for 6, 12 or 24 hours. Higher and lower donor lifetime correspond to higher and lower tension, respectively. **(D)** (Left) ER-Src cells expressing the Vin-TSΔtail and treated with EtOH or TAM for 6, 12 or 24 hours. Color bar indicates 0–6 ns. (Right) Quantification of the mTFP1 lifetime at FAs in ER-Src cells expressing the Vin-TSΔtail and treated with EtOH or TAM for 6, 12 or 24 hours. **(E)** (Left) Confocal images of ER-Src cells treated with EtOH or TAM for 6, or 12 hours, stained with Phalloidin (magenta), anti-Zyxin (yellow) and anti-p120ctn (cyan). (Right) Quantification of FA area in ER-Src cells treated with EtOH or TAM for 6 or 12 hours. Quantifications are from three biological replicates and are presented as violin plots with the magenta lines indicating median values. Statistical significance was calculated using Mann – Whitney t-tests. ns indicates non-significant. *P<0.05; **P<0.01; ****P<0.0001. Scale bars represent 20 μm.

High tension across Vinculin has been associated with FA enlargement (9). Thus, to confirm that FAs are under higher tension in premalignant ER-Src cells, we evaluated the size of Zyxin-positive area 6 and 12 hours after EtOH or TAM treatment (Fig. 2E). Quantifications indicated that the average area of Zyxin puncta was significantly larger in ER-Src cells treated with TAM for 12 hours, compared to those treated with EtOH for the same time (Fig. 2E), suggesting that ER-Src cells assemble larger FAs 12 hours after TAM treatment. Together, we conclude that FAs are under higher tension in premalignant ER-Src, prior cells undergoing EMT.

### Tension at FAs is required to strengthen forces at AJs in premalignant ER-Src cells

Forces on FAs can be transferred to AJs (11). We therefore asked if preventing the transient increase in tension at FAs suppressed the ability of premalignant ER-Src cells to increase tension at AJs. We first tested the effect of inhibiting the activity of FAK, as FAK plays a key role in building up tensile forces at FAs (24). Consistent with previous report (21), FAK phosphorylation (pFAK) was strongly potentiated after 12 hours of treatment with TAM (Supp. Fig. 3A). Blocking FAK phosphorylation with PF-573228, a known FAK inhibitor (FAKi), strongly reduced pFAK levels in cells treated with either EtOH or TAM for 12 hours, compared to those treated with DMSO (Supp. Fig. 3A). Inhibiting FAK activity did not prevent the assembly of FAs, seen by the presence of Zyxin and Vin-TS-positive foci in ER-Src cells co-treated with FAKi and EtOH or TAM (Fig. 3A). However, FAK activity is required for strengthening tensile forces at FAs in premalignant cells, as the extended Vin-TS donor lifetime at FAs 12 hours after TAM treatment was suppressed in the presence of FAKi. In contrast, FAKi did not affect the Vin-TS donor lifetime in EtOH-treated cells (Fig. 3B).

**Figure 3:**
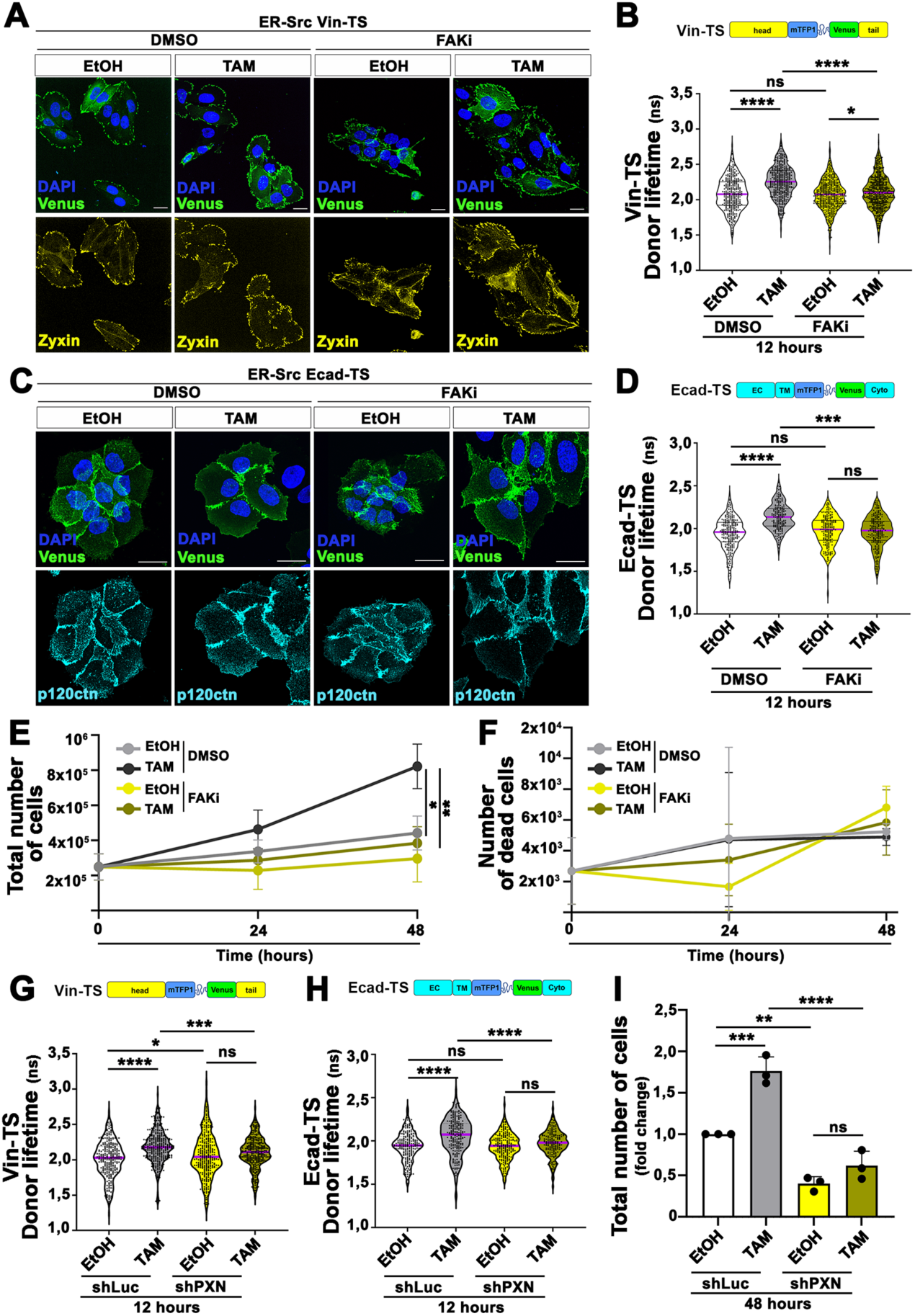
TAM-treated ER-Src cells lose the ability to increase tensile forces at AJs and to grow when unable to strengthen forces at FAs. **(A)** Confocal images of ER-Src cells expressing the Vin-TS (Venus in green) and co-treated with EtOH or TAM and DMSO or FAKi for 12 hours. Cells are stained with anti-Zyxin (yellow) and DAPI (blue). **(B)** Quantification of the mTFP1 lifetime at FAs in ER-Src cells expressing the Vin-TS and co-treated with EtOH or TAM and DMSO or FAKi for 12 hours. **(C)** Confocal images of ER-Src cells expressing the Ecad-TS (Venus in green) and co-treated with EtOH or TAM and DMSO or FAKi for 12 hours. Cells are stained with anti-p120ctn (cyan) and DAPI (blue). **(D)** Quantification of the mTFP1 lifetime at AJs in ER-Src cells expressing the Ecad-TS and co-treated with EtOH or TAM and DMSO or FAKi for 12 hours. Quantifications of donor lifetime are presented as violin plots with the magenta lines indicating median values. Scale bars represent 20 μm**. (E)** Growth curve of ER-Src cells co-treated with EtOH or TAM and DMSO or FAKi during 48 hours. **(F)** Quantification of dead ER-Src cells co-treated with EtOH or TAM and DMSO or FAKi at 24h and 48h timepoints. **(G)** Quantification of the mTFP1 lifetime at FAs in ER-Src cells expressing the Vin-TS and shLuc or shPXN and treated with EtOH or TAM for 12 hours. **(H)** Quantification of the mTFP1 lifetime at AJs in ER-Src cells expressing the Ecad-TS and shLuc or shPXN and treated with EtOH or TAM for 12 hours. **(I)** Quantification of the total number of ER-Src cells expressing shLuc or shPXN and treated with EtOH or TAM for 48 hours. Quantifications are from three biological replicates. Statistical significance was calculated using one-way ANOVA with Tukey’s multiple comparison. ns indicate non-significant. *P<0,05; **P<0.01; ***P<0.001; ****P<0.0001.

We then tested if inhibiting FAK activity affected tension at AJs. ER-Src cells co-treated with FAKi and EtOH or TAM for 12 hours still assembled in clusters with p120ctn and the Ecad-TS localized at the interface between neighboring cells (Fig. 3C). However, FAKi prevented the extended Ecad-TS donor lifetime at AJs in TAM-treated ER-Src cells at 12 hours (Fig. 3D). Moreover, FAKi significantly suppressed the growth of TAM-treated cells at 48 hours (Fig. 3E), but had no major effect on cell viability (Fig. 3F), indicating that FAK activation provides a proliferative advantage to TAM-treated ER-Src cells.

To confirm that increased tension at FAs strengthens forces at AJs and is required for cellular transformation, we knocked down PXN using small-hairpin RNAs (shPXN), as Paxillin is one of the consensus FA components, not shared between AJs and FAs (11,25). EtOH or TAM-treated ER-Src cells expressing shPXN showed reduced Paxillin protein levels, compared to those infected with small-hairpin RNAs against Luciferase (shLuc), used as control (Supp. Fig. 3B). Knocking down PXN in ER-Src cells did not prevent the assembly of Zyxin- and Vin-TS-positive FAs (Supp. Fig. 3C). It did not either induce the loss of cell-cell contacts, as Ecad was still localized between adjacent PXN-depleted cells or between PXN-depleted cells and those expressing Paxillin (Supp. Fig. 3D). However, knocking down PXN suppressed the extended donor lifetime of the Vin-TS (Fig. 3G) and of the Ecad-TS (Fig. 3H) in cells treated with TAM for 12 hours. Moreover, PXN-depleted cells showed a growth impairment, compared to those expressing shLuc (Fig. 3I). Thus, we conclude that the transient strengthening of FAs in premalignant ER-Src cells is required to increase tension at AJs and to sustain cell growth.

### Premalignant ER-Src cells unable to assemble AJs fail to strengthen tension at FAs

We then tested if, in turn, tension at AJs was transferred to FAs in premalignant ER-Src cells. To this end, we disrupted cell-cell adhesion using the calcium chelator EGTA, which prevents *trans* junctional homophilic interactions between cadherin extracellular domains (26). Thirty minutes of EGTA treatment was sufficient to induce cell rounding and the loss of cell-cell contacts in ER-Src cells (Supp. Fig. 4A). After removal of EGTA from the media, cells remained viable for at least 24 hours (Supp. Fig. 4B), and recovered a cobblestone-like shape, with p120ctn localized at cell-cell contacts (Supp. Fig. 4A). This shows that the EGTA-dependent loss of cell-cell contacts is reversible. Thus, we first tested if EGTA affected the size of FAs 12 hours after TAM treatment. As expected, ER-Src cells co-treated with EGTA and EtOH or TAM for 12 hours maintained a rounder shape with E-cad poorly recruited at cell-cell contacts, compared to those grown in the absence of EGTA (Fig. 4A). Quantification of the size of Zyxin-positive puncta indicates that EGTA treatment significantly reduced the average area of Zyxin-positive FAs in both EtOH- and TAM-treated cells (Fig. 4B). Thus, tension at AJs could be required to strengthen forces on FAs. Accordingly, FLIM-FRET on the Vin-TS indicates that the extended donor lifetime 12 hours after TAM treatment was suppressed in the presence of EGTA. In contrast, EGTA had no major effect on the Vin-TS donor lifetime in EtOH-treated cells (Fig. 4C). EGTA treatment also prevented the transient increase in cell stiffening 6 hours after TAM treatment (Fig. 4D). Taken together, these observations suggest that higher tension exerted between TAM-treated ER-Src cells through cadherins, reinforces cortical tension and increases tensile forces at FAs.

**Figure 4:**
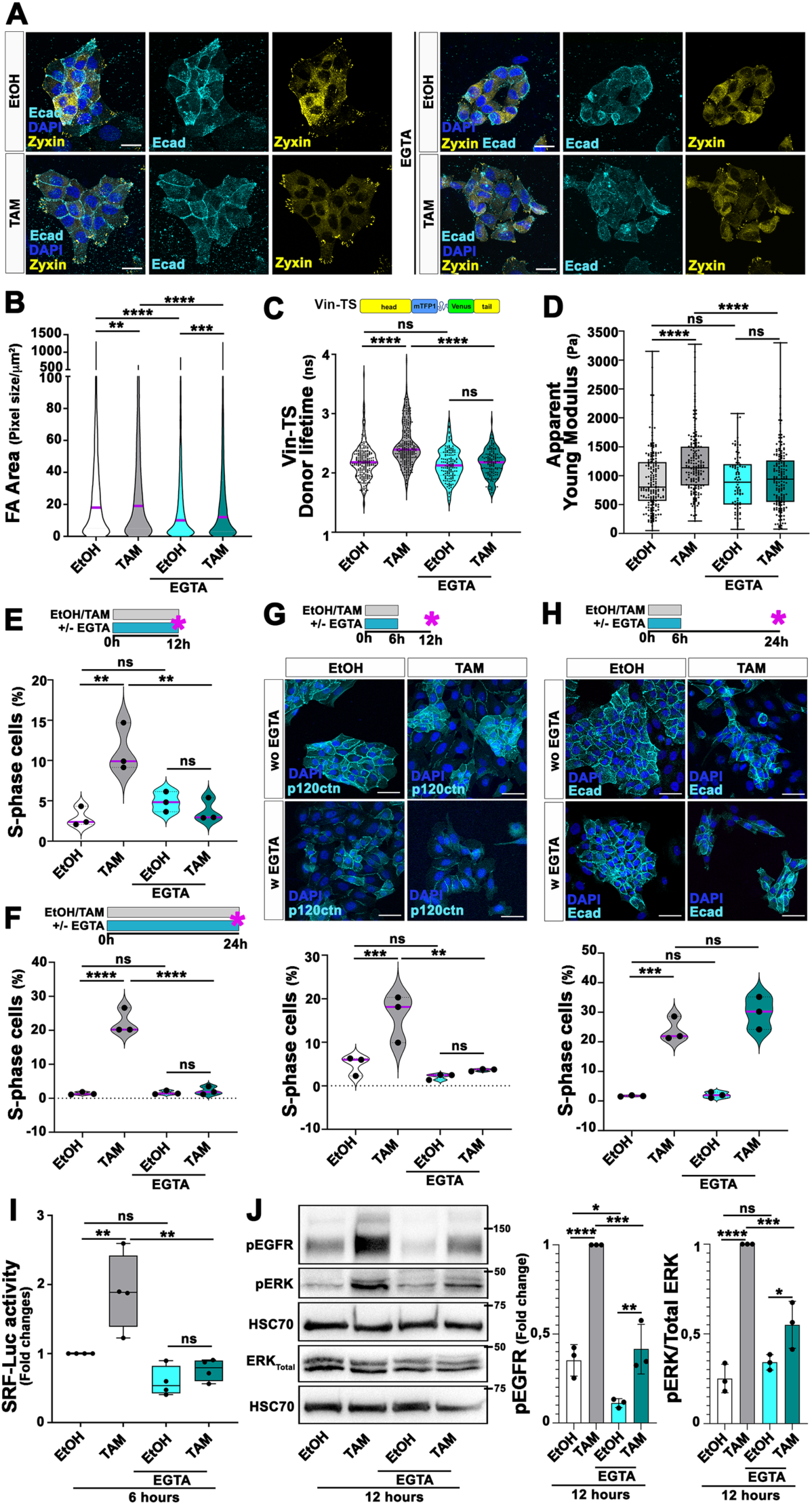
EGTA treatment hampers the ability of TAM-treated ER-Src cells to increase tension at FAs and to acquire sustained proliferative ability. (**A**) Confocal images of ER-Src cells treated with EtOH or TAM for 12 hours in the presence or absence of EGTA, stained with anti-Ecad (cyan), anti-Zyxin (yellow) and DAPI (blue). **(B)** Quantification of FA area in ER-Src cells treated with EtOH or TAM for 12 hours in the presence or absence of EGTA. **(C)** Quantification of the mTFP1 lifetime at FAs in ER-Src cells expressing the Vin-TS and treated with EtOH or TAM for 12 hours in the presence or absence of EGTA. **(D)** Quantification of the Apparent Young’s modulus of ER-Src cells treated with EtOH or TAM for 6 to 8 hours in the presence or absence of EGTA. (**E-H)** (Top) Schematics of the experimental designs. Magenta stars indicate the time points at which cells were collected. (Bottom) Quantification of the percentage of ER-Src cells in S-phase, in cells treated with EtOH or TAM hours in the presence or absence of EGTA for **(E)** 12 hours or **(F)** 24 hours. **(G-H)** (Middle) confocal images or (lower) quantifications of the percentage of cells in S-phase in ER-Src cells treated with EtOH or TAM for 6 hours in the presence or absence of EGTA and left to grow for an additional 6 **(G)** or 18 **(H)** hours in the absence of treatment. Cells are stained with Ecad (cyan) and DAPI (blue). **(I)** Quantification of the fold changes in Luciferase activity in ER-Src cells transfected with the SRF-responsive-Luc reporter gene and treated with EtOH or TAM for 6 hours in the presence or absence of EGTA. **(J)** (Left) Western blots on protein extracts from ER-Src cells treated with EtOH or TAM for 12 hours in the presence or absence of EGTA, blotted with anti-pEGFR, anti-pERK and anti-HSC70 used as loading control (middle) or anti-ERK (ERK_Total_) and anti-HSC70 (bottom). (Right) Quantifications of pEGFR levels or the ratio of pERK over total ERK levels, normalized to HSC70 for the conditions indicated. Quantifications are from three (B to H and J) or four (I) biological replicates and presented as violin plots with the magenta lines indicating median values (B, C and E to H) or as column plots (D, I and J). Statistical significance was calculated using a linear mixed model. ns indicate non-significant. *<0.05; **P<0.01; ***P<0.001; ****P<0.0001. Scale bars indicate 20 μm.

### Premalignant ER-Src cells unable to assemble AJs fail to sustain proliferation

We then tested the consequences of preventing AJ assembly on the ability of TAM-treated ER-Src cells to gain sustained proliferative abilities. Consistent with previous reports (19,20), TAM-treated ER-Src cells showed a significant higher number of cells in S-phase of the cell cycle, 12 (Fig. 4E) and 24 (Fig. 4F) hours after treatment, compared to those treated with EtOH for the same time. However, in the presence of EGTA, TAM-treated cells lost their ability to gain self-sufficiency in growth properties, 12 (Fig. 4E) or 24 (Fig. 4F) hours after Src activation. This effect is not due to cell death induced by EGTA, as EGTA-treated cells survived equally well in all experimental conditions (Supp. Fig. 4C, D). EGTA does not suppress the ability of TAM-treated cells to acquire sustained proliferative abilities by preventing Src activation, as the phosphorylation levels of the ER-Src fusion protein (pER-Src) were not significantly different between cells treated with TAM only and those co-treated with TAM and EGTA (Supp. Fig. 4E).

We then tested if TAM-treated ER-Src cells regain a proliferative advantage after withdrawing EGTA and allowing AJs re-assembly. Cells co-treated with EGTA and TAM or EtOH for 6 hours and left to grow in the absence of treatments for another 6 hours did not recover an epithelial-like morphology with membrane-associated p120ctn at cell-cell contacts. In this experimental setting, EGTA still impeded TAM-treated cells to gain proliferative competency (Fig. 4G), with no effect on cell survival (Supp. Fig. 4F). However, 18 hours after withdrawing EGTA and EtOH, cells completely restored an epithelial-like morphology and those that received EGTA and TAM survived equally well and regained the ability to enter S-phase (Fig. 4H and Supp. Fig. 4G).

We then analyzed the impact of EGTA treatment on MRTF-A-SRF and ERK activities, as both pathways provide a proliferative advantage to premalignant ER-Src cells (19,20). As previously reported (20), the activity of a Luciferase reporter controlled by three SRF binding sites (SRF-Luc) was significantly higher in ER-Src cells 6 hours after TAM treatment, compared to those treated with EtOH. Consistent with a role of cell-cell adhesion in enabling premalignant ER-Src cells to gain proliferative abilities, EGTA suppressed the transient increase in SRF-dependent Luciferase activity in TAM-treated cells (Fig. 4I). EGTA also significantly suppressed the higher phosphorylation levels of EGFR (pEGFR) and ERK (pERK) 12 hours after TAM treatment (Fig. 4J). In these cells, ERK phosphorylation is dependent on EGFR activation, as co-treating TAM-treated ER-Src cells with the EGFR inhibitor tyrphostin AG1478 (EGFRi) strongly reduced pEGFR and pERK levels in both EtOH- and TAM-treated cells (Supp. Fig. 4H). Taken together, these observations suggest that higher tensile forces at AJs provide a proliferative advantage to premalignant ER-Src cells through the activation of the EGFR-ERK and MRTF-A-SRF signaling pathways.

### P-cadherin builds up tension at AJs and FAs in premalignant ER-Src cells

If higher tensile forces exerted between premalignant ER-Src cells through cadherin homophilic interactions are required to strengthen tension at FAs and for cellular transformation, interfering with cadherin function should phenocopy the behavior of EGTA-treated cells. MCF10A cells express the two classical cadherins Ecad and Pcad (27). According to a role of Pcad-mediated AJ strengthening in cellular transformation, we reported that the transient accumulation of membrane-associated Pcad in premalignant ER-Src cells promotes a boost of MRTF-A-SRF activity, which sustains cell proliferation (20). We therefore asked if Pcad builds up tension at AJs and FAs in premalignant ER-Src cells. ER-Src cells transfected with a small interference RNA against Pcad (siPcad) showed a striking decrease of membrane-associated Pcad, compared to those expressing control small interference RNAs (siCTR). Yet, these cells assembled in clusters with Ecad localized at the interface between neighboring cells (Fig. 5A). Pcad is required to build up tension at AJs in premalignant cells, as the extended donor lifetime of the Ecad-TS in ER-Src treated with TAM for 12 hours was significantly suppressed by expressing siPcad, compared to those expressing siCTR (Fig. 5B). Knocking down Pcad also prevented the transient extended lifetime of the Vin-TS at FAs in ER-Src cells 12 hours after TAM treatment (Fig. 5C). In contrast, reducing Pcad function in EtOH-treated cells slightly increased the donor lifetime of the Ecad-TS (Fig. 5B) and Vin-TS (Fig. 5C). We then analyzed the effect of Pcad on the activation of EGFR-ERK signaling. ER-Src cells treated with TAM for 12 hours and expressing siPcad displayed a strong reduction in Pcad, pEGFR and pERK levels, compared to those expressing siCTR (Fig. 5D).

**Figure 5:**
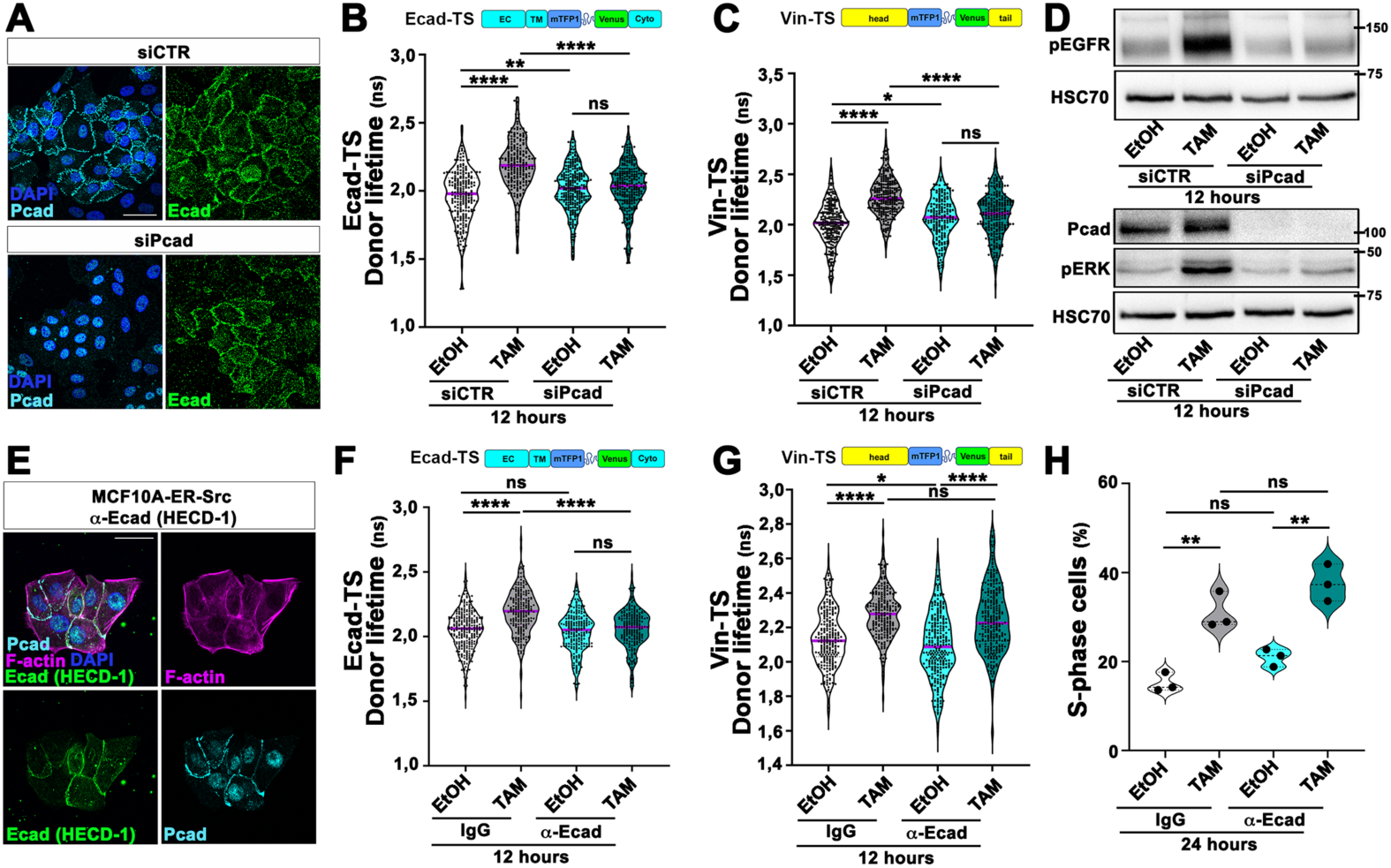
Pcad, but not Ecad, increases tensile forces at AJs and FAs. **(A)** Confocal images of ER-Src cells expressing siCTR or siPcad, stained with anti-Pcad (cyan), anti-Ecad (green) and DAPI (blue). **(B)** Quantification of the mTFP1 lifetime at AJs in ER-Src cells expressing the Ecad-TS and siCTR or siPcad and treated with EtOH or TAM for 12 hours. **(C)** Quantification of the mTFP1 lifetime at FAs in ER-Src cells expressing the Vin-TS and siCTR or siPcad and treated with EtOH or TAM for 12 hours. **(D)** Western blots on protein extracts from ER-Src cells expressing siCTR or siPcad and treated with EtOH or TAM for 12 hours, blotted with anti-pEGFR and anti-HSC70 used as loading control or anti-Pcad, anti-pERK and anti-HSC70. **(E)** Confocal images of ER-Src cells incubated with 10 μg/mL of the Ecad blocking antibody HECD-1 (green), stained with Phalloidin (magenta), anti-Pcad (cyan) and DAPI (blue). **(F)** Quantification of the mTFP1 lifetime at AJs in ER-Src cells expressing the Ecad-TS and treated with EtOH or TAM in the presence of IgG or Ecad blocking antibody for 12 hours. **(G)** Quantification of the mTFP1 lifetime at FAs in ER-Src cells expressing the Vin-TS and treated with EtOH or TAM in the presence of IgG or Ecad blocking antibody for 12 hours. **(H)** Quantifications of the percentage of ER-Src cells in S-phase treated with EtOH or TAM for 24 hours in the presence of IgG or anti-Ecad blocking antibody. Quantifications are from three biological replicates and are presented as violin plots with the magenta lines indicating median values. Statistical significance was calculated using one-way ANOVA with Tukey’s multiple comparison. ns indicate non-significant. *P<0,05; **P<0.01; ***P<0.001; ****P<0.0001. Scale bar indicates 20 μM.

We then tested the role of Ecad in increasing tensile forces at FAs and in cellular transformation. Unlike Pcad, the levels of the mature and pro-peptide Ecad forms were not significantly different between cells treated with EtOH and TAM for 12 hours (Supp. Fig, 5). To analyze the consequence of affecting Ecad function on FA strengthening, we blocked the *trans* junctional homophilic interactions between the Ecad extracellular domains using the Ecad blocking antibody HECD-1. As expected, HECD-1 was found associated with the cell membrane. These cells were still assembled in clusters with Pcad localized at the interface between neighboring cells (Fig. 5E). Although the Ecad blocking antibody prevented the extended donor lifetime of the Ecad-TS in cells treated with TAM for 12 hours (Fig. 5F), it did not suppress the higher Vin-TS donor lifetime in these cells (Fig. 5G), nor their ability to enter S-phase (Fig. 5H). Taken together, we conclude that higher tensile forces at AJs, exerted between TAM-treated ER-Src cells through Pcad homophilic interactions, is required for building up tension at FAs and for ERK-dependent proliferation.

### Cadherin is required for Src-induced overgrowth *in vivo*

We then tested if AJs were required for Src-induced overgrowth *in vivo*. *Drosophila* Ecad (DEcad), encoded by *shotgun*, is the main classical cadherin contributing to cell-cell adhesions in the wing disc epithelium (28). We therefore tested if knocking down *DEcad* suppresses the overgrowth of *Drosophila* wing discs overexpressing the *Src oncogene at 64B (Src64B)* and the caspase inhibitor *p35* using the UAS-Gal4 system. As expected, distal wing discs overexpressing *Src64B, p35* and two copies of Green Fluorescent Protein (GFP) with the *nubbin-*Gal4 (*nub*-Gal4) driver were bigger than control discs expressing GFP only. Strikingly, replacing one copy of UAS-GFP by a UAS construct expressing double-strand RNA (dsRNA) directed against *Ecad (DEcad-IR#1)* appeared to reduce the growth of *Src/p35*-expressing wing discs (Fig. 6A). Quantification of the ratio between the GFP-positive area and the total wing area for each experimental condition confirmed that knocking down *DEcad* significantly suppressed the overgrowth of *Src/p35*-expressing wing discs (Fig. 6A). Expressing an independent dsRNA against *DEcad (DEcad-IR#2)* also significantly suppressed the overgrowth of *Src64B/p35*-expressing wing discs (Fig. 6B). To confirm that the overgrowth suppression observed reflected indeed a reduction in cell number, we also evaluated the percentage of GFP-positive cells by cell sorting for each experimental condition. While the number of GFP-positive cells was significantly increased by 22% in *nub>Src64B, p35-*expressing wing discs, compared to control discs expressing *GFP* only, knocking down *DEcad* in these tissues restored the percentage of GFP-positive cells to those of control discs expressing *GFP* only (Fig. 6C and Supp. Fig. 6). These observations are in agreement with a role for AJs in Src-induced tissue growth.

**Figure 6:**
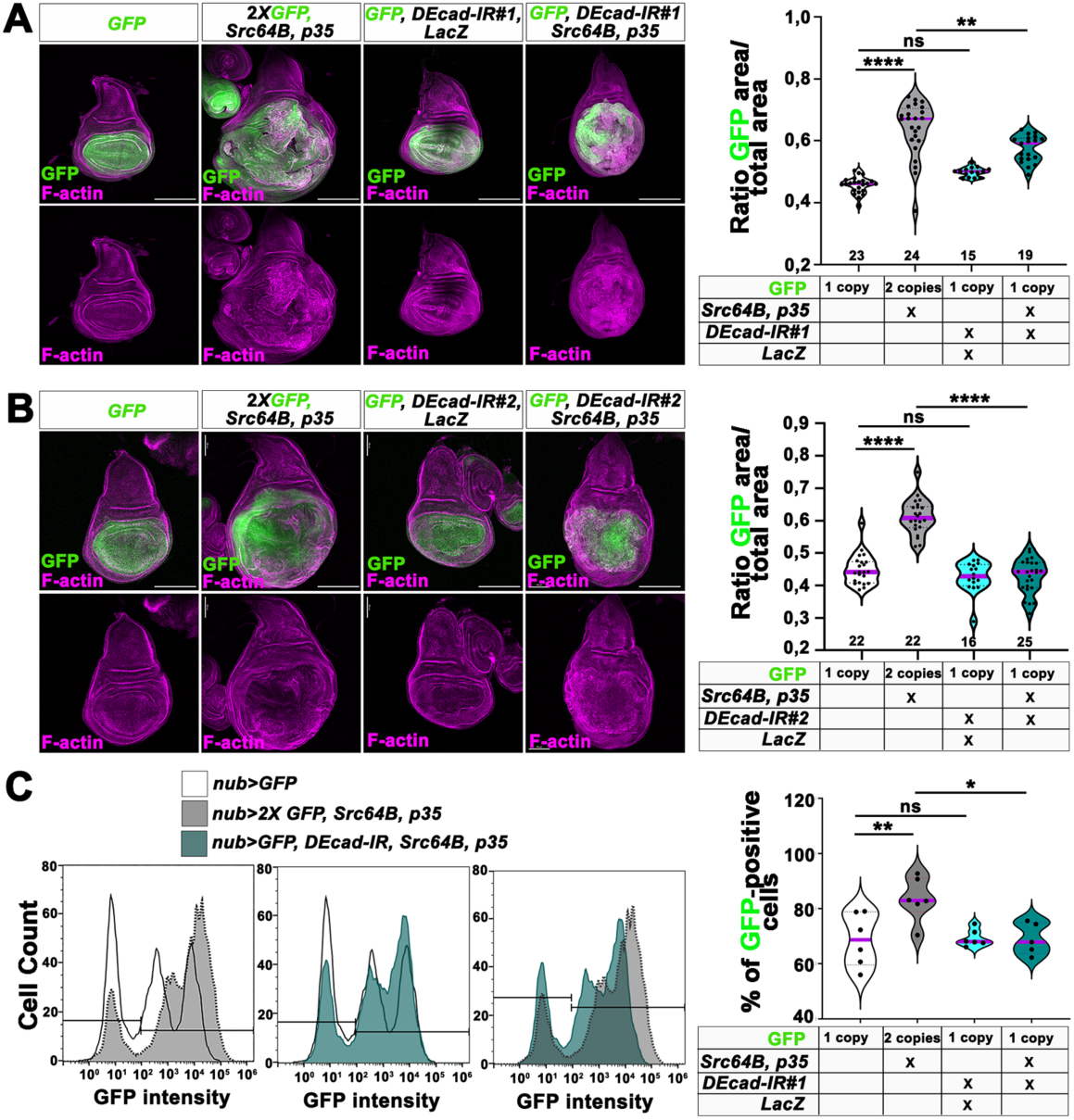
Knocking down *DEcad* suppresses the overgrowth induced by Src overexpression *in vivo*. **(A)** (Left) Standard confocal sections of third instar wing imaginal discs with dorsal side up, expressing one copy of UAS-CD8::GFP (green) or two copies of UAS-*CD8::GFP* (green), UAS-*p35* and *Src64B^UY1332^* or one copy of UAS-*CD8::GFP* (green), UAS-*LacZ* and UAS-*DEcad-IR^GL00646^* or one copy of UAS-*CD8::GFP* (green), UAS-*DEcad-IR^GL00646^*, UAS-*p35* and *Src64B^UY1332^* under *nub-*Gal4 control. Discs are stained with Phalloidin (magenta). (Right) Ratio of the *nub>GFP* area over the total wing disc area for the genotypes indicated. Number of samples analyzed are indicated on the X axis, and are from two biological replicates. **(B)** (Left) Standard confocal sections of third instar wing imaginal discs with dorsal side up, expressing one copy of UAS-CD8::GFP (green) or two copies of UAS-*CD8::GFP* (green), UAS-*p35* and *Src64B^UY1332^* or one copy of UAS-*CD8::GFP* (green), UAS-*LacZ* and UAS-*DEcad-IR^GD14421^* or one copy of UAS-*CD8::GFP* (green), UAS-*DEcad-IR^GD14421^*, UAS-*p35* and *Src64B^UY1332^*under *nub-*Gal4 control. Discs are stained with Phalloidin (magenta). (Right) Ratio of the *nub>GFP* area over the total wing disc area for the genotypes indicated. Number of samples indicated on the X axis are from three biological replicates. **(C)** (Left) Histograms of membrane-associated GFP levels in wing discs from the genotypes indicated. (Right) Percentage of GFP-positive cells in wing discs from the genotypes indicated. Quantifications are from five biological replicates. Quantifications are presented as violin plots with the magenta lines indicating median values. Statistical significance was calculated using one-way ANOVA with Tukey’s multiple comparison. ns indicate non-significant. **P<0.01; ***P<0.001; ****P<0.0001. Scale bar indicates 20 μM.

## Discussion

The weakening of cell-cell and cell-matrix adhesiveness has been proposed to allow cancer cells to acquire hallmarks of malignant tumors, including sustaining proliferative signaling, resisting cell death and activating invasion and metastasis (29). Here, we provide evidence that AJs and FAs have a paradoxical role in premalignant breast epithelial cells overactivating Src. We show that prior to undergoing EMT, a transient mechanical crosstalk between Pcad-mediated forces transmission at AJs and FAs provide a proliferative advantage to ER-Src cells through activation of the EGFR-ERK and MRTF-A-SRF signaling (Fig. 7).

**Figure 7:**
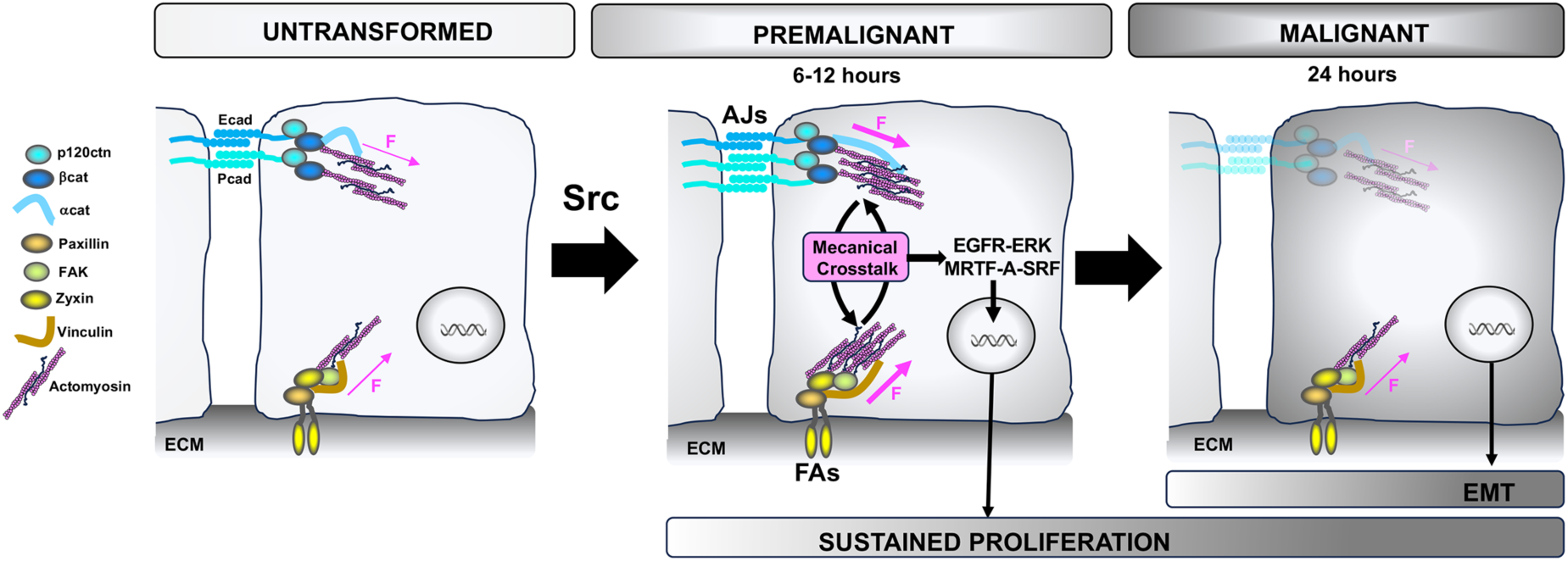
Model by which the transient mechanical crosstalk between AJs and FAs in premalignant ER-Src cells sustains cell proliferation, prior to undergoing EMT. Premalignant Src-overactivating MCF10A cells accumulate membrane-associated Pcad, which increases tensile forces at AJs and FAs. In turn, FAs transmit tensile forces to AJs. This positive mechanical crosstalk triggers activation of the EGFR-ERK and MRTF-A-SRF signaling pathways, which provide to TAM-treated ER-Src cells a proliferative advantage. Later, cells lose membrane-associated Ecad and Pcad, reduce tensile forces at FAs and acquire features associated with EMT.

Our observations demonstrate that prior to triggering features associated with EMT, Src transiently buildups tension at AJs and FAs that are transferred between both adhesion sites. During the first 12 hours of Src activation, Ecad and αcat at AJs and Vinculin at FAs bear higher forces that correlate with the assembly of larger FAs. Moreover, premalignant cells unable to assemble AJs due EGTA or that fail to strengthen tension at AJs due to Pcad depletion, lose the ability to reinforce tension at FAs. Conversely, preventing the strengthening of tensile forces at FAs by restricting FAK or Paxillin activity, suppresses the ability of premalignant ER-Src cells to build up higher tensile forces at AJs. The requirement for increased tensile forces at AJs prior to EMT may not be unique to MCF10A cells with conditional Src activation. In the *Drosophila* wing disc epithelium, tension at AJs is required in pre-tumoral tissues for subsequent tumor evolution (30). Likewise, Hepatocyte Growth Factor (HGF) enhances homophilic Ecad interaction in MDCK cells prior to inducing AJ disassembly (31). EGF also increases the magnitude of tensile force on Ecad in MCF10A cells prior to triggering cell scattering (32). Thus, major alterations in AJs and FAs could constitute prime events in cancer development, ahead of any signs of EMT. Consistent with this possibility, we reported that the two most significant pathways affected in premalignant breast atypical hyperplasia are components of the AJ and FA adhesomes (19).

We propose that higher tensile stresses at AJs provide a proliferative advantage to Src-overactivating cells by promoting activation of the EGFR-ERK and MRTF-A-SRF signaling pathways. Consistent with this model, TAM-treated ER-Src cells unable to assemble AJs due to EGTA, lose the ability to gain a proliferative advantage and to enhance EGFR-ERK and MRTF-A-SRF signaling activities. In turn, ERK and MRTF-A are required to sustain proliferation in those cells (19,20). Moreover, TAM-treated cells regain self-sufficiency in growth capacity if empowered to re-assemble AJs. Furthermore, *in vivo*, DEcad, the sole classical cadherin involved in AJ-dependent cell-cell adhesions, enables the overgrowth of Src-overexpressing wing discs. In agreement with our observations, Myosin II activity is required for ERK-mediated proliferation in premalignant ER-Src cells (19). Mechanical stresses can also activate EGFR-ERK and trigger MRTF-A nuclear translocation in other cell models (33,34). Ligand-independent ERK activation can be induced by Ecad extracellular engagement, which phosphorylates EGFR at cell-cell contact in keratinocytes (35). Premalignant ER-Src cells do not require Ecad trans-homophilic interaction to enter S-phase. Rather, this function is endorsed by Pcad, which transiently accumulates at the cell membrane and triggers a boost of MRTF-A-SRF signaling activity (20). In addition, we show that Pcad is required to potentiate EGFR-ERK activity in premalignant cells. Pcad also triggers EGFR signaling in oral dysplastic keratinocytes (36). Moreover, in premalignant ER-Src cells, Pcad is necessary to increase tensile forces on Ecad at AJs. Consistent with our observations, Pcad mediates intercellular force transmission in MCF10A cells (37). However, in breast cancer cell lines, Pcad reduces tension at AJs and destabilizes the interaction between Ecad and catenins (38,39). Pcad overexpression is also strongly associated with cytoplasmic p120ctn accumulation in pancreatic and ovarian cancer cell lines (40,41). Thus, Pcad may have a dual opposite effect on AJ strengths that could depend on the stage of tumor progression and therefore on the composition of the AJ adhesomes. Yet, the mechanism by which the Pcad-dependent force transmission activates the EGFR-ERK and MRTF-A-SRF signaling pathways remains to be characterized.

In addition, to provide a proliferative advantage to premalignant ER-Src cells, EGFR activation mediated by force transmission at AJs may also build up tensile stresses at FAs. In agreement with this model, EGFR activation increases tension on integrin and controls the spatial organization of FAs in Cos-7 cells (42). Likewise, in MCF7 cells, EGFR activation increases FAs size and number and integrin-mediated traction forces (17). Yet, we cannot exclude that Pcad-mediated force transmission affects EGFR-ERK- and MRTF-A-SRF activities through increasing tensile forces at FAs. EGFR has also been shown to interact with integrin in human fibroblasts and endothelial cells and is activated by integrin-mediated adhesions even in the absence of EGF, leading to ERK activation (43). In turn, EGFR signaling can reinforce AJs and junctional actomyosin, as shown in the *Drosophila* ectoderm prior to gastrulation onset (44). Integrin-mediated force transmission can also trigger MRTF-A nuclear translocation (45). Moreover, we show that FAK and Paxillin are required for force transmission at FAs in premalignant ER-Src cells, as well as for sustaining cell proliferation. However, as FAK and Paxillin also strengthen tension at AJs, the growth defect of Src-overactivating ER-Src cells defective for FAK or Paxillin activity could result from the shortfall in AJ strengthening. Forces transmitted from FAs to AJs could further increase tensile forces at AJs, which, upon reaching a certain threshold may trigger the subsequent dissociation of F-actin from AJs and the loss of cell-cell adhesions required for EMT, as proposed in endothelial cells (46). Thus, while uncontrolled growth because of the loss of cell-cell contacts has been proposed to enable the initiation and progression of several types of cancer (47), our work reveals that a transient mechanical crosstalk between AJs and FAs in premalignant breast cells could drive tumor growth.

## Methods

### Fly strains and genetics

Fly stocks used were UAS-*DEcad-IR^GD14421^* (Vienna Drosophila Research Center, VDRC), and *nub-*Gal4, Src64B^UY1332^, UAS-*p35*, UAS-*DEcad-IR^GL00646^* (Bloomington Drosophila Stock Center). All crosses were maintained at 25 °C. Male and female larvae were dissected at the end of the third instar.

### Cell culture conditions and treatments

The MCF10A-ER-Src cell line (ER-Src for short), kindly provided by K. Struhl (48), was grown in Complete Growth Media (CGM), composed of DMEM/F12 growth media (Gibco, 11039-047), supplemented with 5% of Charcoal Stripped Horse Serum (CSHS) (Gibco, 16050-122), 20 ng/mL human EGF (Peprotech, AF-100-15), 0.5 µg/mL Hydrocortisone (Sigma, H0888), 100 ng/mL Cholera toxin from Vibrio cholerae (Sigma, C8052), 10 µg/mL Insulin (Sigma, I9278), 0.5 µg/mL Puromycin (Merck, 540411) and 1% Penicilin-Streptomycin (Gibco, 15140). To treat cells with 1 µM 4OH-TAM (Merck, H7904) or EtOH, cells were plated in CGM at 50% confluency and allowed to adhere for 12 to 24 hours. Cells were then treated with 1 µM 4OH-TAM (Merck, H7904) or an equal volume of EtOH for the time indicated To analyze the effect of EGTA, cells were plated in CGM at 50% confluency, allowed to adhere for 12 to 24 hours and starved in Restricted Growth Media (RGM), composed of DMEM/F12, supplemented with 0.5% CSHS, 20 ng/mL human EGF, 0.5 µg/mL hydrocortisone, 100 ng/mL cholera toxin, 10 µg/mL insulin, 0.5 µg/mL Puromycin and 1% Penicillin-Streptomycin for 12 hours. Cells were then treated with 4 mM EGTA (BioWorld, 40520008) or left without treatment for 30 minutes, and co-treated with 1 µM 4OH-TAM or an equal volume of EtOH for the indicated time. For cell cycle analysis, RGM media without EGF was used.

To block the *trans* junctional homophilic interactions between Ecad extracellular domain, cells were seeded and allowed to adhere in the presence of 10 µg/mL of anti-Ecad monoclonal antibody (HECD-1, Invitrogen, 13-1700) or with an identical volume of IgG in CGM. Cells were then starved for 12 hours in RGM lacking EGF in the presence of HECD-1 or IgG and co-treated with 1 µM 4OH-TAM or EtOH and HECD-1 or IgG for 24 hours for cell cycle analysis or directly co-treated with 1 µM 4OH-TAM or EtOH and HECD-1 or IgG in CGM for all other experiments for the time indicated.

To inhibit FAK activity, cells were plated in CGM and allowed to adhere for 24 hours. Cells were then treated with 8 µM PF-573228 (Sigma-Aldrich, PZ0117) or DMSO for 12 hours in RGM lacking EGF for cell cycle analysis or for 2 hours in CGM for all other experiments. Cells were then co-treated with EtOH or 1 µM 4OH-TAM and PF-573228 or DMSO for the indicated time in RGM lacking EGF for cell cycle analysis or CGM for all other experiments. To inhibit EGFR activity, cells were seeded in CGM and allowed to adhere for 24 hours. Then, cells were washed twice, grown in DMEM/F12 for 12 hours, and treated with 2 µM of the EGFR inhibitor for 1 hour, and co-treated with 1 µM 4OH-TAM or EtOH in the presence or absence of 2 µM AG1478 (Santa Cruz Biotechnology, sc-200613) for the indicated time.

The Human Embryonic Kidney 293T cells (HEK293T) used for viral production were grown in DMEM/F12 supplemented with 10% Fetal Bovine Serum (FBS) Premium heat-inactivated (Thermo Fisher Scientific, 10500-064). MDA-MB-231 cells were grown in Dulbecco’s Modified Eagle Medium (DMEM) High Glucose 1X + GlutaMAX™-I (Thermo Fisher Scientific, 31966), supplemented with 10% FBS and 1% Penicillin-Streptomycin (Thermo Fisher Scientific, 15070-063).

All cell lines were grown in a humidified incubator at 37 °C, under a 5% CO_2_ atmosphere. Tests for mycoplasma contamination were performed every other month, via PCR using the primers MGS0 5′-TGCACCATGTGTCACTCTGTTAACCTC-3′ and GPO1 5′-ACTCCTACGGGAGGCAGCAGTA-3′, which amplify the 16S ribosomal RNA genes of the Mollicutes class of mycoplasma.

### Molecular Biology

Ecad-TS and Ecad-TSΔCyto constructs were generated from the canine EcadTSMod sequence kindly provided by Alex Dunn lab (Stanford University) (10). Briefly, the cassette containing the Ecad-TS or Ecad-TSΔCyto was excised with EcoRI and XbaI, and inserted within the pGenLenti vector. Vin-TS and Vin-TSΔtail constructs were generated from the Addgene plasmids #26019 and #26020, kindly provided by Martin Schwartz (9). Both plasmids were excised from their backbone using the HindIII and SacII restriction enzymes and inserted into the pGenLenti vector.

### Lentivirus production and infection

To produce shLuc (Sigma Aldrich, TRCN0000123137), shPXN (Sigma Aldrich, TRCN0000123137), Ecad-TS, Ecad-TSΔcyto, Vin-TS, or Vin-TSΔtail viral particles, HEK293T cells were seeded to reach 70-80% confluency and transfected with 6 μg of pMD2.G (envelope vector), 17 μg of psPAX2 (packaging vector), and 22 μg of plasmids of interest using Lipofectamine 2000 (Thermo Scientific, 11668019). Supernatants were collected 48 and 72 hours after transfection, centrifuged at 500 g for 10 minutes, and filtered using a 0.45 μm filter unit to remove large cellular debris, followed by concentration through ultracentrifugation using the Optima L-80 XP Ultracentrifuge (Beckman Coulter) at 32,000 *g* for 1.5 hours at 4°C. Pellets were resuspended in CGM, aliquoted, and stored at -80°C. To infect ER-Src cells, plated in 6-well plates, CGM was replaced with 1 mL of CGM supplemented with 10 μg/mL Polybrene® (EMD Millipore, TR-1003-G). Lentiviral particles were added dropwise to each well and mixed by gentle swirling. 12h later, 1 mL of CGM media was added to each well. Cells were incubated for 72 hours and assessed for protein expression by flow cytometry or western blot.

### FRET-FLIM Analysis

To perform FRET-FLIM experiments, ER-Src cells stably expressing each FRET-based sensor (Ecad-TS, Ecad-TSΔCyto, Vin-TS, Vin-TSΔtail) were plated on an ibiTreat 8-well chamber slide (Ibidi, 80826) in CGM and allowed to adhere for 24h. After treatment, images were acquired on a single point scanning confocal Stellaris 8, equipped with a fully motorized inverted Leica DMI8 microscope, an HC PL APO 86x/1.20 W motCORR STED white water immersion objective equipped with a correction collar (Leica Microsystems), a WLL, and Falcon and STED modalities, using the Leica LASX software (version 4.6.1.27508). Cells were selected using the acceptor fluorescence signal (EYFP) to ensure the proper localization of the tension sensor (AJs for Ecad-TS and FAs for Vin-TS). The Navigator function on LASX was used to have an overview of the sample and to identify clusters of cells. Fluorescent proteins were excited using a white light laser (WLL) at 85% power, tuned to 80 MHz. The WLL was tuned to 440 or 514 nm to excite mTFP1 and EYFP, respectively, with laser powers of 2% and 2%. Photon detection was performed using two Power HyD X detectors in photon counting (intensity mode), tuned to 440 nm (mTFP1). Images were recorded sequentially by frame scanning unidirectionally at 400 Hz using the galvanometer-based imaging mode, with a line accumulation of 16, and a pixel size of 0.176 μm for an area size of 20,83 μm x 20,83 μm (512 x 512, zoom factor of 1, pixel dwell time of 3.1625 μs). For each experiment, a minimum of 20 cell clusters were analyzed.

FRET detection was based on the time-domain FLIM experiments, which were performed using time-correlated single-photon counting. At least 100 photon events per pixel were collected in all cases, and the lifetime analysis was performed using the Phasor approach on LASX.

### Immunofluorescence Analysis

Wing imaginal discs from third instar larvae were dissected in a phosphate buffer at pH 7 (0.1 M Na2HPO4, 0.1 M NaH2PO4 at a 72:28 ratio). Discs were then fixed in 4 % formaldehyde in PEM (0.1 M PIPES pH 7, 0.2 mM MgSO4, 1 mM EGTA) for 20 to 30 min, rinsed in phosphate buffer 0.2 % Triton for 15 minutes and incubated 2 hours at 4°C with Rhodamine-conjugated Phalloidin (Sigma, P-1951) at 0,3 mM in phosphate buffer 0.2 % Triton supplemented with 10 % horse serum. Wing discs were washed 3 times, 10 minutes each, before being mounted in Vectashield (Vector Labs, H-1000). Fluorescence images were acquired on a single point scanning confocal SP5, equipped with a fully motorized inverted Leica DMI6000 microscope, an HC PL APO Lbl. Blue 20x/0.70 IMM objective equipped with a correction collar (Leica Microsystems).

For cells, 1.5×10^5^ ER-Src or 1×10^5^ E-cad TS, Vinc-TS, Ecad-TSΔcyt, Vin-TSΔtail cells were seeded on poly-L-lysine-coated coverslips and allowed to adhere for 12 to 24 hours. After treatment, cells were rinsed with PBS 1X and fixed with 4% formaldehyde methanol-free (Thermo Fisher Scientific, 28908) in H_2_O, supplemented with 0.06 M PIPES pH 6.8, 0.03 M HEPES pH 7.0, 0.01 M EGTA pH 6.8, and 0.004 M MgSO_4_ for 10 minutes. Cells were permeabilized with PBS containing 0.1% Triton-X-100 (PBS-T) for 2 minutes at RT and blocked with a blocking buffer (10% FBS in PBS-T) for 1 hour at RT. Rat anti-α18 (1:50; a gift from Professor Akira Nafaguchi, Japan, (23), mouse anti-p120ctn (1:50; BD Bioscience, 610133), rabbit anti-Zyxin (1:50; Sigma, HPA004835), mouse anti-Ecad (1:50; Invitrogen, 13-1700), mouse anti-Pcad (1:50; BD Bioscience, 610228), rabbit anti-Ecad (1:50; Cell Signalling, 3195) and mouse anti-Paxillin (1:50 or 1:200; BD Bioscience, 610051) were incubated overnight at 4°C in blocking buffer. Coverslips were washed with PBS 1X three times for 5 minutes at RT and incubated with FITC-conjugated Donkey anti-Rat IgG (1:200; Jackson ImmunoResearch, 712-095-150), FITC-conjugated Donkey anti-Mouse IgG (1:200; Jackson ImmunoResearch, 715-095-150), Alexa Fluor 647-conjugated Donkey anti-Mouse IgG (1:200; Jackson ImmunoResearch, 715-605-151), CyTM5-conjugated Donkey anti-Rabbit IgG (1:200; Jackson ImmunoResearch, 711-175-152), and Rhodamine-conjugated Phalloidin (Sigma-Aldrich, P-1951) at 0.3 mM in blocking buffer for 1 hour at RT in the dark. After three washes with PBS 1X for 5 minutes, cells were stained with DAPI (1:500; Sigma, D9542) for 5 minutes at RT, washed with PBS 1X for 5 minutes, and mounted in Vectashield. Images were obtained on a single-point scanning confocal Leica SP5, coupled to a Leica DMI6000, using the 63x/1.4 HCX PL APO CS oil immersion objective or with a spinning disk confocal Andor BC43 benchtop system (Oxford Instruments/Andor) using the 60x/1.42 oil immersion objective. Image processing was performed using Imaris version 10.2 or NIH Fiji/ImageJ.

### Immunoblotting Analysis and Quantification

2.65×10^5^ ER-Src cells in CGM were plated in a T25 flask. After 6 or 12 hours of treatment, cells were trypsinized with TrypLE^TM^ Express (Gibco, 12604-021) and collected into 1.5 mL tubes. Proteins were extracted by resuspending the cell pellets in Lysis Buffer SDS-Free, composed of 0.05M Tris-HCl pH 7.5 (VWR, 33621.260), 0.15 M NaCl (Sigma-Aldrich, 31434), 0.001 M EDTA pH 8.0 (Sigma-Aldrich, E5134), 0.001 M EGTA pH 7.0 (Sigma-Aldrich, E3889), 1% Triton X-100 (Sigma-Aldrich, T8787) and supplemented with 1% protease inhibitors (Sigma-Aldrich, 5892791001) and 1% phosphatase inhibitors (Sigma-Aldrich, 4906845001) on ice for 30 minutes. Samples were centrifuged at 4°C for 30 minutes at 14,000 rpm, and protein was quantified using the Bradford method. For pERK and pEGFR detection, 1.5×10^5^ ER-Src cells were plated in 6-well plates. After 12 hours of treatment, Lysis Buffer SDS-Free, containing 1% protease inhibitors (Sigma-Aldrich, 5892791001) and 1% phosphatase inhibitors (Sigma-Aldrich, 4906845001; Sigma-Aldrich, S6508; Sigma-Aldrich, S7920), was added to cells. Cells were detached using a cell scraper and collected into 1.5 mL tubes. Samples were centrifuged at 4°C for 30 minutes at 14,000 rpm, and protein was quantified using the Bradford method. Laemmli buffer was added to protein samples to a final concentration of 1X, and extracts were boiled for 5 minutes at 95°C, centrifuged for 10 minutes at 8,000 rpm before being resolved by SDS-PAGE electrophoresis and transferred to a 0.45 µM PVDF blotting membrane (Amersham, 10600023). Membranes were blocked with 5% BSA in TBS containing 0.1% Tween 20 for 1 hour, cut to separate proteins migrating at diverse molecular weights and incubated with rat anti-α18 (1:1,000; (Nagafuchi & Tsukita, 1994)), rabbit anti-αcat (1:1,000; Thermo Fisher Scientific, MA5-14986), mouse anti-HSC70 (1:8,000; Santa Cruz Biotechnology, sc-7298), mouse anti-Paxillin (1:10,000; BD Bioscience, 610051), mouse anti-phospho-p44/42 MAPK (1:1,000; Cell Signaling, 9106), rabbit anti-p44/42 MAPK (1:2,000; Cell Signaling, 9102), rabbit anti-GAPDH (1:5,000; Sigma, G9545), rabbit anti-pEGFR (1:1,000; Thermo Fisher Scientific, 44-788G) mouse anti-Pcad (1:2,500; BD Bioscience, 610228), rabbit anti-pFAK (pY397) (1:1,000; Cell Signalling), rabbit anti-Ecad (1:1,000; Cell Signalling, 3195) and rabbit anti-phospho Src (pY418) (1:3,000; Invitrogen, 44-660G) diluted in TBS 0.1% Tween 20 supplemented with 3% BSA. Secondary antibodies used were horseradish peroxidase (HRP)-conjugated AffiniPure donkey anti-mouse IgG (1:5,000; Jackson ImmunoResearch, 715-035-150), HRP-conjugated AffiniPure donkey anti-rabbit IgG (1:5,000; Jackson ImmunoResearch, 715-035-152), goat anti-rat IgG-HRP (1:5,000; Santa Cruz biotechnology, sc-2006), IRDye® 680RD-conjugated donkey anti-mouse IgG (1:20,000; LI-COR Bioscences, 926-68072) and IRDye® 800CW-conjugated donkey anti rabbit IgG (1:20,000; LI-COR Bioscences, 926-32213), all diluted in TBS 0.1% Tween 20 supplemented with 1% BSA. Immobilon Western Chemiluminescent HRP substrate (Millipore, P90719) was utilized for detection and visualization in ChemiDoc (Bio-Rad) or Odyssey® M (LI-COR Biosciences) imaging systems. Image Lab (Bio-Rad) and Image Studio (LICOR) software were used for quantification. The intensity levels of each band were normalized to those of the respective control for protein extracts loaded on the same gel for each experimental condition.

### Transfections and dual Luciferase Renilla reporter assay

To test the effect of knocking down Pcad, 1.5×10^5^ ER-Src cells resuspended in 2 mL of CGM were plated in 6-well plates for immunoblot analysis or in ibiTreat 8-well chamber slide for FRET-FLIM analysis. After 12 to 24 hours, cells were transfected with 50 nM or 100nM siCTR (Qiagen, 1027310) or 50 nM or 100nM siCDH3 (Qiagen, 1027416, GeneGlobe ID:SI02663941) for FRET-FLIM and immunofluorescence or for immunoblot analysis, respectively, using Lipofectamine 2000, according to the manufacturer’s instructions. After 24 hours, transfected cells were incubated for another 12 hours in CGM containing EtOH or 4OH-TAM for FRET-FLIM or immunofluorescence analysis. For immunoblot analysis, 12 hours after transfection, the medium was replaced by RGM lacking EGF. After 36 hours, cells were treated with 4OH-TAM or EtOH for the indicated time. To test the effect of EGTA on the activity of the SRF-Luc reporter, 1×10^5^ cells resuspended in 2 mL of CGM were plated in 6-well plates. 12 to 24 hours later, cells were transfected with 0.720µg p3D.A-Luc and 1.44 µg pRL-TK plasmids (49) using Lipofectamine 2000, according to the manufacturer’s instructions. Cells were then incubated with EGTA for 30 minutes, and co-treated with 4OH-TAM or EtOH in the presence or absence of EGTA for 6 hours in RGM lacking EGF. After trypsinization with TrypLE™ Express (Thermo Fisher Scientific, 12604021), cell pellets were resuspended in 250 µL 1× passive lysis buffer (Promega, E1941) and stored overnight at −20°C. Luciferase assay was performed using a Dual-Luciferase Reporter Assay system (Promega, PROME19600010) according to the manufacturer’s instructions. Briefly, 100 µL Luciferase assay reagent II (LARII) was added to 80 µL of lysed samples in a 96-well plate. Firefly luminescence was detected using Synergy Mx (BioTek). Then, 100 µL 1X Stop and Glo solution (Promega, PROME19600010) was added to detect Renilla luminescence. Each sample was evaluated in triplicate.

### Cell Cycle Profile

For cell cycle analysis, cells treated for 12 or 24 hours were trypsinized with TrypLE^TM^ Express and collected into 5 mL round-bottom polystyrene tubes (Corning, 352235). Cells were centrifuged for 5 minutes at 1,000 rpm at 4°C, washed with 1 mL of FACS Buffer (PBS 1X supplemented with 2% heat-inactivated FBS (Biowest, S181BH)), centrifuged for 5 minutes at 1,000 rpm at 4°C and resuspended in 500 µL FACS Buffer. Cells were fixed with 1.5 mL 70% Ethanol, added drop-by-drop, while gently vortexed, followed by a 30 minutes incubation at 4°C. After 5 minutes centrifugation at 2,000 rpm at 4°C, cells were resuspended in 3 mL of ice-cold PBS 1X, incubated 30 minutes on ice, pelleted again by centrifugation for 5 minutes at 2,000 rpm at 4°C and resuspended in 300 µL of PBS 1X, supplemented with 100 µg/mL RNAse A (Qiagen, 19101) and 20 µg/mL Propidium Iodide (Sigma, P4170). Samples were incubated for 30 minutes in a 37°C water bath in the dark. Flow Cytometry was performed at a low flow rate using a BD Accuri C6 Flow Cytometer (Becton-Dickinson, Franklin Lakes, NJ, USA). Cell Cycle profiles were analyzed using FlowJo 10.7.1 software (Tree Star, Inc., Ashland, OR, USA) Cell Cycle platform, using the Watson Model.

### Quantification of cell number

To quantify cell number, cells were trypsinized with TrypLE^TM^ Express, washed twice with PBS 1X, and resuspended in PBS 1X. Then, cell suspension was mixed with trypan blue in a 1:1 proportion, pipetted into a disposable Countess cell counting chamber slide (Thermo Fisher Scientific, C10228) and cell number was quantified using the Countess 3 Automated Cell Counter (Thermo Fisher Scientific).

### Flow Cytometry

To quantify GFP-expressing cells in wing imaginal discs, 25 discs were dissected for each genotype in ice-cold PBS 1X. Samples were then incubated in 200 µL of TrypLE™ for 45 minutes at 37°C to facilitate cell dissociation. Mechanical pipetting was performed to ensure complete dissociation before inactivating the TrypLE™ with 200 µL of PBS 1X supplemented with 1% FBS. Cells were centrifuged at 3,500 rpm for 8 minutes, fixed in 500 µL of ice-cold 70 % Ethanol, and stored at 4°C. Before analysis, cells were resuspended in 400 µL of PBS 1X, and filtered into 5 mL FACS round-bottom tubes. 100 µL of cell suspension was analyzed with medium flow rate settings on a BD Accuri C6 Plus. A description of the gating strategy can be found in Supp. Fig. 6. To confirm the expression of the FRET-tension sensors, transduced ER-Src cells were washed and resuspended in PBS 1X supplemented with 0.2% BSA and 0.1% NaN3, at a concentration of 1×10^6^ cells/ml. Cytometric analysis was performed using a FACSCanto II or LSRFortessa flow cytometers (BD), with FACS Diva software.

### Atomic force microscopy

1×10^5^ cells in 2 mL of CGM were plated in a glass bottom dish (World Precision Instruments, FD35-100). 6 hours after treatment with EtOH or TAM in the absence or presence of EGTA, RGM was replaced by CO_2_-independent media (Thermo Fisher Scientific 18045054) with EtOH or 4OH-TAM in the presence or absence of EGTA. EGTA induced the appearance of two morphologically distinct cell populations in both EtOH- and TAM-treated cells. The round cell populations were excluded from the analysis, as we assume that these cells are losing adherence to the glass substrate. For the AFM indentation experiments, a Nanowizard 4 (JPK Instruments, Berlin) mounted on top of a light microscope (Zeiss Axiobserver) was used, combined with a petri dish heater to keep the temperature at 37°C during experiments. Arrow-T1 cantilevers (Nanoworld, Neuchatel, Switzerland) were modified with polystyrene beads (radius 2.5 μm, Microparticles GmbH, Berlin, Germany) using epoxy glue to obtain a well-defined spherical indenter geometry and decrease local strain during indentation. The cantilever was lowered over the central part of the cell with a speed of 5 μm s^-1^ until a relative set point of 2.5 nN was reached. Each cell was probed three times in the same location. The resulting force-distance curves were analyzed using JPK image processing software (JPK instruments). Force-distance data were corrected for the tip sample separation and fitted with a Hertz model for a spherical indenter to extract the apparent Young’s Modulus within an indentation depth of approximately 1 μm. A Poisson ratio of 0.5 was assumed. The average of the Young’s Modulus of the three curves was calculated for each cell. A 2-hour time window was required for measurement of the four experimental conditions.

### Quantifications

The NIH Fiji/ImageJ program was used to quantify the ratio of the GFP-positive wing disc area to the total disc area. Comparisons between wing imaginal discs overexpressing Src64B and p35 alone or together with UAS-Ecad-IR were performed using the same number of UAS transgenes. For each wing disc, the area of the GFP domain and the area of the whole disc domain highlighted by Phalloidin staining were outlined manually and measured using the *Area* function, which evaluates size in square pixels. The ratio was calculated for each disc between the two. To quantify the intensity of the α18 staining, maximum intensity Z-projections of the p120ctn and α18 channels were performed for each field using the NIH Fiji/ImageJ program. The segmented line tool was used to overlap the p120ctn signal with the line width set to 10. Each line was then used to measure the mean gray value in the α18 channel. Three fields per condition were quantified for each biological replicate. To quantify FA size, maximum intensity Z-projections of the Zyxin channel were performed for each field using the NIH Fiji/ImageJ program. Images were then segmented using the LABKIT Fiji plugin to generate binary masks. The size of particles was then extracted and displayed in pixel units. To compare the size of FAs between ER-Src cells treated with EtOH or TAM for 6 or 12 hours, images were acquired on a single point scanning confocal SP5, equipped with a fully motorized inverted Leica DMI6000 microscope (Leica Microsystems), with a 63x objective using a zoom factor 3 to analyze FA area between ER-Src cells treated with EtOH or TAM for 6 or 12 hours or a zoom factor 2 to determine the effect of EGTA on the FA area. 3 to 5 fields per condition were quantified for each biological replicate. To analyze FA area between ER-Src cells treated with EtOH or TAM for 6 or 12 hours, particles smaller than 20 pixels were considered nonspecific signals and were excluded. To analyze the effect of EGTA on the FA area, all particles were included. R was used for the statistical analysis of AFM data. GraphPad Prism (10.0) was used for all other statistical analysis and for data presentation.

## Acknowledgements

We specially thank K. Struhl, Alex Dunn, G. Posern, A. Nagafuchi, A. Dunn and M. Schwartz for reagents and all members of F.J. lab for helpful discussions. We acknowledge the Cell Culture and Genotyping, the Genomics, the Translational Cytometry, the Bioimaging and the Advanced Light Microscopy platforms at i3S. Stocks obtained from the Bloomington Drosophila Stock Center (NIH P40OD018537) and from the Vienna Drosophila Research Center, VDRC, were used in this study. This research was funded by Fundação para a Ciência e Tecnologia I.P. (FCT), grant number PTDC/BIA-BFS/0812/2021 and 2022.06763.CEECIND to F.J. and PhD fellowship number 2022.11122.BD (doi https://doi.org/10.54499/2022.11122.BD) to L.F. The i3S Bioimaging and Advanced Light Microscopy scientific platforms are both members of the national infrastructure PPBI-Portuguese Platform of BioImaging (POCI-01-0145-FEDER-022122).

## Author contributions

Conceptualization F.J.; Methodology F.J., M.A., P.S., A.T.; Investigation, L.F., C.G., I.F.; C.S.L; V.R., S.T., V.M., F.J.; Formal Analysis, L.F., C.G., I.F.; C.S.L; V.R., S.T., V.M., F.J.; Writing – Original Draft, F.J.; Writing – Review & Editing, L.F., C.G., I.F.; C.S.L; V.R., S.T., V.M., F.J.; Funding Acquisition, F.J.; Supervision, F.J., A.T. and L.F.

## Data Availability

Correspondence and material requests should be addressed to.

