## Supplementary Figures for "Prior to Undergoing Epithelial to Mesenchymal Transition, Premalignant Cells Transiently Increase Tensile Forces at Cell-Cell and Cell-Matrix Adhesions"

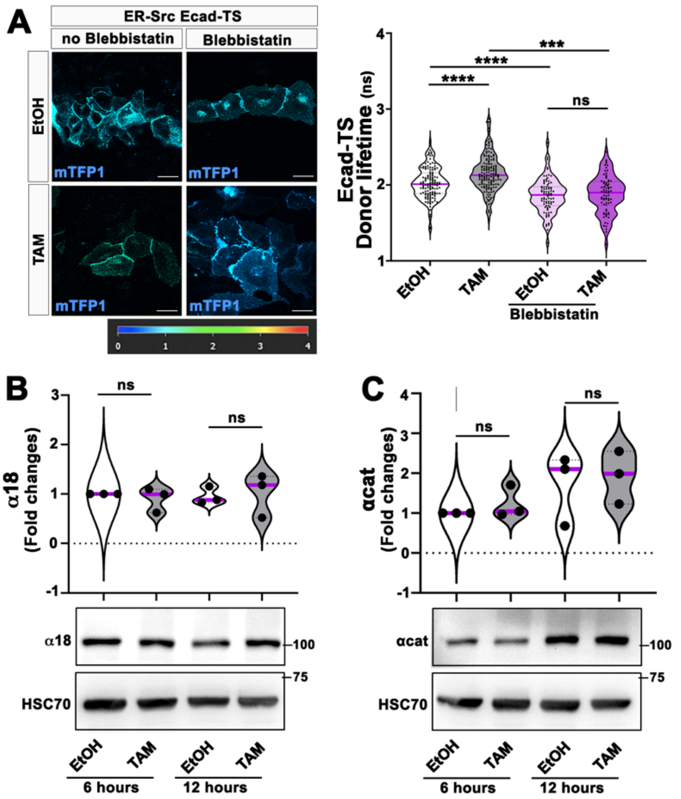


**Supplementary Figure 1: The increased lifetime of the Ecad-TS in TAM-treated ER-Src cells depends on Myosin II but does not involve alteration in αcat levels.**

**(A)** (Left) ER-Src cells expressing the Ecad-TS (mTFP1) and treated with EtOH or TAM in the presence or absence of Blebbistatin for 6 hours. Color bar indicates 0–4 ns. (Right) Quantification from three biological replicates of the mTFP1 lifetime at AJs in ER-Src cells expressing the Ecad-TS and treated with EtOH or TAM in the presence or absence of Blebbistatin for 6 hours.

**(B, C)** (Bottom) Western blots on protein extracts from ER-Src cells treated with EtOH or TAM for 6 or 12 hours, blotted with α18 (B) or anti-αcat (C) and anti-HSC70 (B and C). (Top) Quantifications from three biological replicates of the α18 (B) or αcat intensity (C), normalized to HSC70 for the corresponding lane on western blot.

Quantifications are presented as violin plots with the magenta lines indicating median values. Statistical significance was calculated using one-way ANOVA with Tukey's multiple comparison. ns indicates non-significant. ***$P<0.001$; ****$P<0.0001$. Scale bar indicates 20 µM.

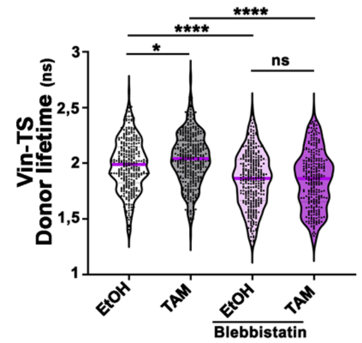


**Supplementary Figure 2: The increased lifetime of the Vin-TS in TAM-treated ER-Src cells depends on Myosin II.**

Quantification from three biological replicates of the mTFP1 lifetime at FAs in ER-Src cells expressing the Vin-TS and treated with EtOH or TAM in the presence or absence of Blebbistatin for 12 hours. Quantification is presented as a violin plot with the magenta lines indicating median values. Statistical significance was calculated using one-way ANOVA with Tukey’s multiple comparison. ns indicates non-significant. *$P<0.05$; ****$P<0.0001$.

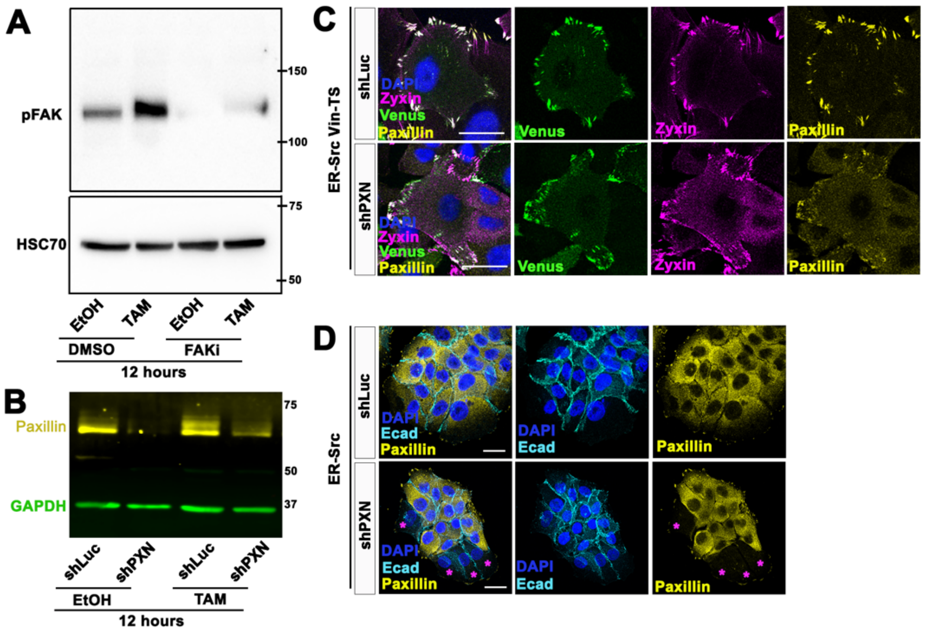


**Supplementary Figure 3: Expressing shPXN in EtOH- or TAM-treated ER-Src cells decreases Paxillin levels but does not prevent FA and AJ assembly, while co-treating ER-Src cells with EtOH or TAM and PF-573228 reduces pFAK levels.**

**(A)** Western blot on protein extracts from ER-Src cells treated with EtOH or TAM and DMSO or PF-573228 (FAKi) for 12 hours, blotted with anti-pFAK or anti-HSC70.

**(B)** Western blots on protein extracts from ER-Src cells expressing shLuc or shPXN and treated with EtOH or TAM for 12 hours, blotted with anti-Paxillin (yellow) and anti-GAPDH (green).

**(C)** Confocal images of ER-Src cells expressing the Vin-TS (Venus in green) and shLuc or shPXN, stained with anti-Zyxin (magenta), anti-Paxillin (yellow) and DAPI (blue). Scale bars indicate 10 µm.

**(D)** Confocal images of ER-Src cells expressing shLuc or shPXN, stained with anti-Ecad (cyan), anti-Paxillin (yellow) and DAPI (blue). Scale bars indicate 20 µm. Magenta asterisks represent ER-Src cells knocked-down for PXN.

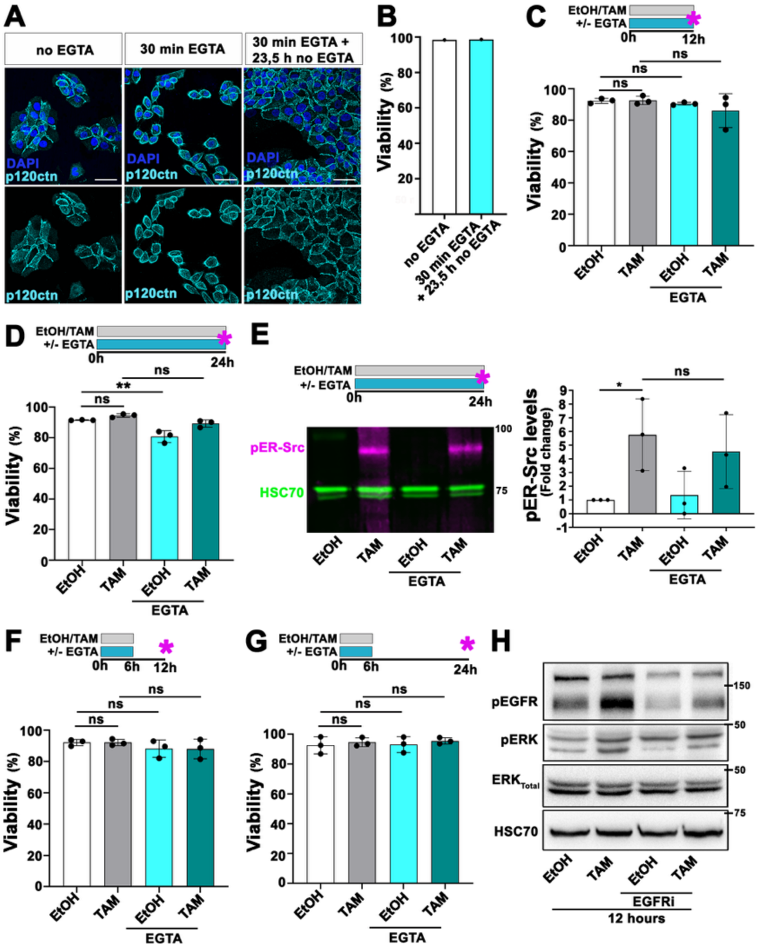


**Supplementary Figure 4: EGTA treatment disrupts cell-cell contacts in ER-Src cells in a reversible manner, without affecting cell viability.**

**(A)** Confocal images of ER-Src cells before or after treatment with 4 mM EGTA for 30 minutes or with 4 mM EGTA for 30 minutes and left to grow for additional 23,5 hours in the absence of EGTA. Cells are stained with anti-p120ctn (cyan) and DAPI (blue).

**(B)** Percentage (%) of viable ER-Src cells untreated or treated with 4 mM EGTA for 30 minutes and left to grow for another 23,5 hours in the absence of EGTA.

**(C-D)** (Top) Schematics of the experimental designs. Magenta stars indicate the time points at which cells were collected for cell cycle analysis. (Bottom) Percentage of viable ER-Src cells treated with EtOH or TAM in the presence or absence of EGTA for **(C)** 12 or **(D)** 24 hours.

**(E)** (Left top) Schematics of the experimental design. Magenta star indicates the time points at which cells were collected for immunoblotting analysis. (Left bottom) Western blots on protein extracts from ER-Src cells treated with EtOH or TAM for 24 hours in the presence or absence of EGTA, blotted with anti-pSrc (magenta) and anti-HSC70 (green). (Right) Quantification from three biological replicates of pER-Src levels, normalized to HSC70 for the corresponding lane on western blot.

**(F-G)** (Top) Schematics of the experimental designs. Magenta stars indicate the time points at which cells were collected for cell cycle analysis. (Bottom) Percentage of viable ER-Src cells treated with EtOH or TAM for 6 hours in the presence or absence of EGTA and left to grow for another **(F)** 6 hours or **(G)** 18 hours in the absence of treatments.

**(H)** Western blots on protein extracts from ER-Src cells treated with EtOH or TAM for 12 hours in the presence or absence of AG1478 (EGFRi), blotted with anti-pEGFR, anti-pERK, anti-ERK ($ERK_{Total}$) and anti-HSC70 used as loading control.

Statistical significance was calculated using one-way ANOVA with Tukey's multiple comparison. Ns indicates non-significant. *P<0,05; **P<0,01. Scale bar indicates 20 um.

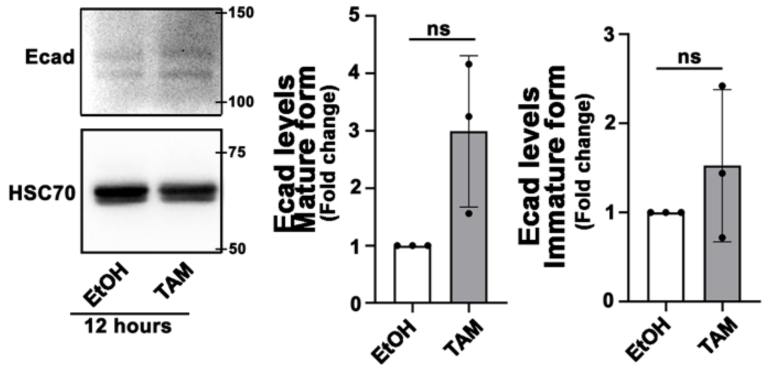


**Supplementary Figure 5: Ecad levels are not significantly affected in premalignant ER-Src cells.**

(Left panels) Western blots on protein extracts from ER-Src cells treated with EtOH or TAM for 12 hours, blotted with anti-Ecad and anti-HSC70. (Middle panel) Quantification from three biological replicates of the levels of the mature Ecad form (lower band), normalized to HSC70, on protein extracts from ER-Src cells treated with EtOH or TAM for 12 hours. (Right panel) Quantification from three biological replicates of the levels of the Ecad pro-peptide (upper band), normalized to HSC70 on protein extracts from ER-Src cells treated with EtOH or TAM for 12 hours. Statistical significance was calculated using one-way ANOVA with Tukey's multiple comparison. ns indicates non-significant.

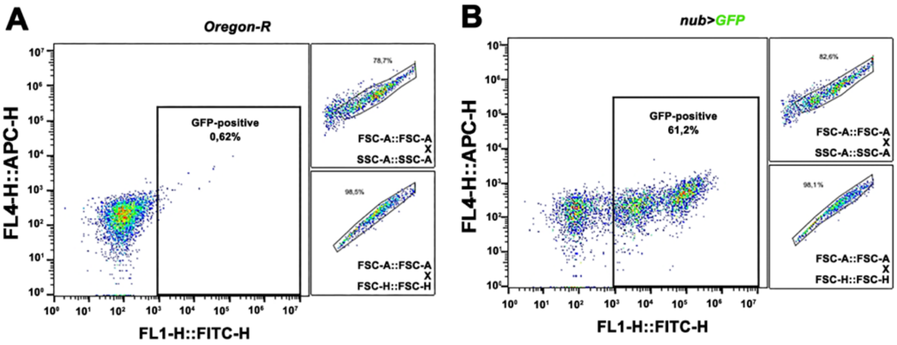


**Supplementary Figure 6: Flow cytometry gating strategy for sorting GFP-positive and GFP-negative cells in *Drosophila* wing discs.**

**(A)** Wild type Oregon wing discs were used as control for GFP-negative cells. (Left) Dot plot of GFP fluorescence shown on the x-axis (log scale). The GFP-positive (0,62%) cells are boxed in black. (Top right) Dot plot of FSC-A versus SSC-A for selection of live GFP-negative cells. (Bottom right) Dot plot of FSC-A versus FSC-H for singletes selection of GFP-negative cells.

**(B)** Wing discs in which *nub*-Gal4 drives the expression of UAS-*CD8::GFP* were used to select GFP-positive cells. (Left) Dot plot of GFP fluorescence shown on the x-axis (log scale). The GFP-positive (61,2%) cells are boxed in black. (Top right) Dot plot of FSC-A versus SSC-A for selection of live GFP-positive cells. (Bottom right) Dot plot of FSC-A versus FSC-H for singlets selection of GFP-positive cells.
